# Transcriptomic analysis of human and mouse muscle during hyperinsulinemia demonstrates insulin receptor downregulation as a mechanism for insulin resistance

**DOI:** 10.1101/556571

**Authors:** Haoning Howard Cen, José Diego Botezelli, Su Wang, Nilou Noursadeghi, Niels Jessen, James A. Timmons, James D. Johnson

**Author notes:** Corresponding author: James D. Johnson, Address: 2350 Health Sciences Mall, Vancouver, BC, Canada, V6T 1Z3; Twitter: @JimJohnsonSci. **Competing Interest Statement:** The authors declare that they have no conflicts of interest with the contents of this article. **Author Contributions:** HC designed the study, performed all experiments except the high-fat feeding, analyzed data, and wrote the manuscript. JDB performed in vivo high-fat feeding and analyzed the related data. SW performed the statistical analysis and edited the manuscript. NN performed siRNA knockdown experiments. NJ contributed to the RNAseq data and edited the manuscript. JT contributed to the design of the transcriptomic analysis and edited the manuscript. JDJ designed the study, supervised the research, and edited the manuscript. JDJ is the ultimate guarantor of this work.

## Abstract

Hyperinsulinemia is commonly viewed as a compensatory response to insulin resistance, yet studies have suggested that chronically elevated insulin may also drive insulin resistance. The molecular mechanisms underpinning this potentially cyclic process remain poorly defined, especially on a transcriptome-wide level. To study the direct effects of prolonged exposure to excess insulin in muscle cells, we incubated C2C12 myotubes with elevated insulin for 16 hours, followed by 6 hours of serum starvation, and established that acute AKT and ERK signaling were attenuated in this model of *in vitro* hyperinsulinemia. Global RNA-sequencing of cells both before and after nutrient withdrawal highlighted genes in the insulin signaling, FOXO signaling, and glucose metabolism pathways indicative of ‘hyperinsulinemia’ and ‘starvation’ programs. We observed that hyperinsulinemia led to a substantial reduction in insulin receptor (*Insr)* gene expression, and subsequently a reduced surface INSR and total INSR protein, both *in vitro* and *in vivo*. Transcriptomic meta-analysis in >450 human samples demonstrated that fasting insulin reliably and negatively correlated with insulin receptor (*INSR*) mRNA in skeletal muscle. Bioinformatic modeling combined with RNAi, identified SIN3A as a negative regulator of *Insr* mRNA (and JUND, MAX, and MXI as positive regulators of *Irs2* mRNA). Together, our analysis identifies novel mechanisms which may explain the cyclic processes underlying hyperinsulinemia-induced insulin resistance in muscle, a process directly relevant to the etiology and disease progression of type 2 diabetes.

## Introduction

Hyperinsulinemia and insulin resistance are cardinal features of type 2 diabetes (T2D) yet their co-association makes it challenging to establish their precise molecular interactions. Insulin resistance has been widely viewed as the primary cause of T2D and hyperinsulinemia is, therefore, a purely compensatory response (1, 2). However, a growing body of evidence suggests the opposite may be true in many cases (3-5). Hyperinsulinemia can be observed prior to insulin resistance in obesity and T2D (6-8). Increased insulin precedes increased BMI (9) and is associated with future T2D in longitudinal studies (10, 11). We recently used a loss-of-function genetic approach to directly demonstrate that hyperinsulinemia can cause age-dependent insulin resistance in the absence of hyperglycemia (12). Reducing hyperinsulinemia in partial insulin gene knockout mice also prevents and/or reverses diet-induced obesity in adult mice (12-14). Rodents (15, 16), healthy humans (17, 18). Further, people with type 1 diabetes (19) subjected to prolonged insulin administration have reduced insulin responsiveness independent of hyperglycemia, strongly implying that intermittent hyperinsulinemia can self-perpetuate or cause insulin resistance.

The mechanisms by which hyperinsulinemia would drive insulin resistance remain poorly understood, particularly at a genome-wide level. Insulin signaling regulates the expression of numerous genes (20) through kinase signaling cascades that culminate in transcription factors (21). Euglycemic-hyperinsulinemic clamp studies have identified genes regulated during acute (∼3h) insulin infusion *in vivo* (22). However, the multiple interacting and time-dependent effects of the hyperinsulinemic clamp on systematic metabolism make it challenging to identify the direct and lasting effects of elevated insulin *in vivo*. Cell culture provides a more constrained model for isolating the primary effects of hyperinsulinemia. An early study by de Campillo et al (23) of the time-dependent transcriptomic responses in muscle cells to physiological insulin (20 nM) identified strong feedback to gene expression in the canonical insulin signaling pathways, yet no impact on the expression of the insulin receptor. It remains unclear how hyperinsulinemia-induced insulin resistance in a cell model impacts the transcriptome, and whether such changes closely mimic *in vivo* observations.

In the present study, we characterized a muscle cell model of hyperinsulinemia-induced insulin resistance and establish that the transcriptomic changes in our *in vitro* model are consistent with those observed in human skeletal muscle across a range of insulin resistant states. We further identify transcriptional regulators that play important roles in mediating the effects of hyperinsulinemia, illuminating how hyperinsulinemia contributes to insulin resistance.

## Materials and Methods

### Cell culture

The C2C12 mouse myoblast (ATCC cell line provided by Dr. Brian Rodrigues, University of British Columbia, Vancouver, Canada) was maintained in Dulbecco’s modified Eagle’s medium (DMEM, Invitrogen) supplemented with 10% (v/v) fetal bovine serum (FBS, Gibco), and 1% penicillin-streptomycin (100 μg/ml; Gibco). For downstream analysis, 8□×□10^5^ cells/well of cells were seeded in 6-well plates and cultured at 37 °C under 5% CO_2_. Confluent (90%) myoblasts were differentiated into myotubes by culturing the cells in differentiation medium (DMEM supplemented with 2% horse serum and 1% penicillin-streptomycin) for 10 days. To induce insulin resistance by hyperinsulinemia *in vitro*, C2C12 myotubes were cultured in the differentiation medium containing 2 or 200 nM human insulin (Cat.# I9278, Sigma) for 16 hours prior to reaching day 10 (Fig.1A). Insulin concentrations after the 16 h hyperinsulinemia treatment were determined using human insulin RIA kit (Millipore). To mimic starvation, myotubes were maintained in the serum-free medium (DMEM supplemented with 1% penicillin-streptomycin) for 6 hours prior to harvesting. All experiments were repeated with biological replicates using cells from different passages.

**Figure 1.**
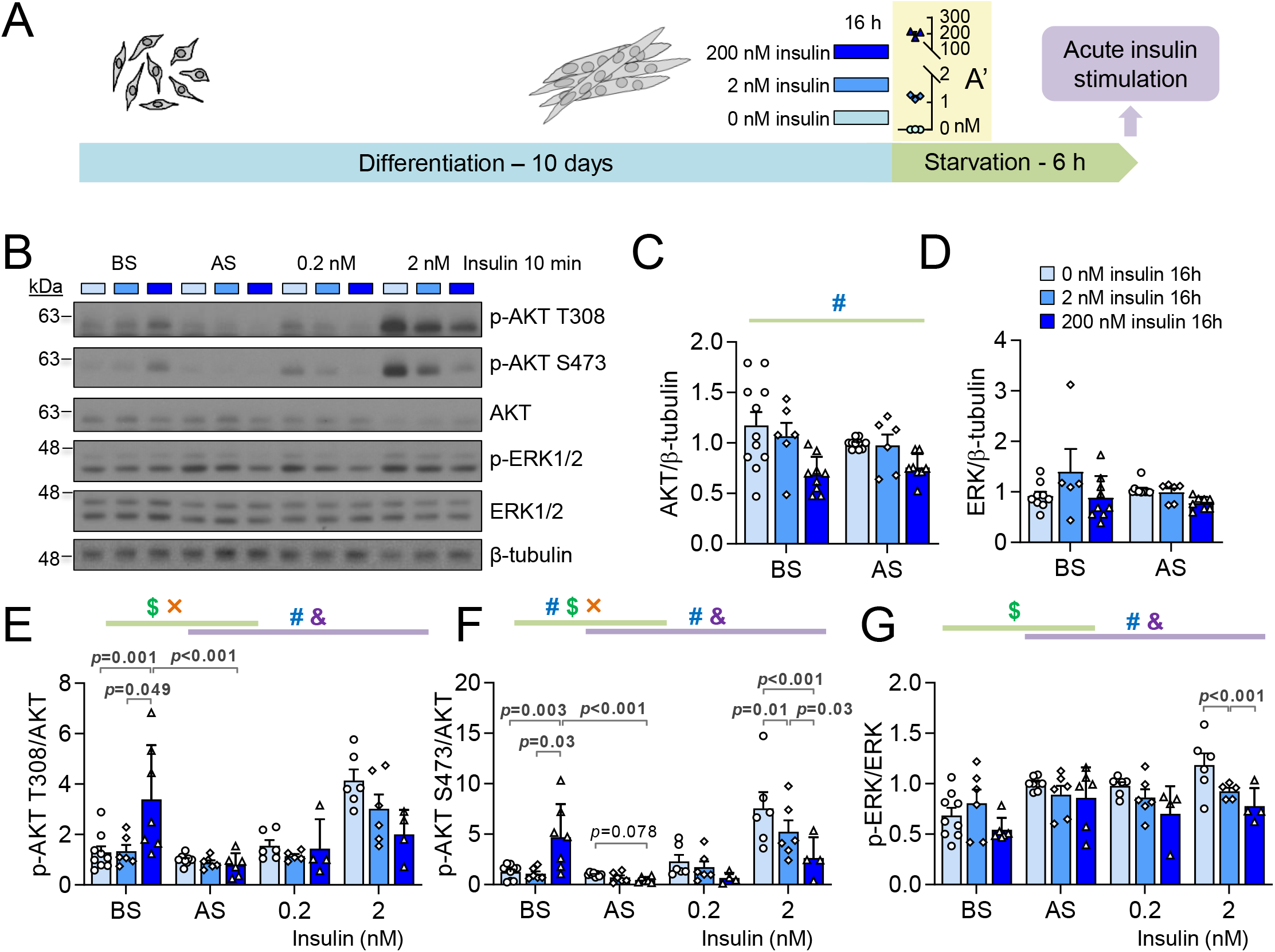
Basal and acute insulin signaling in an *in vitro* hyperinsulinemia-induced insulin resistance model. **(A)** The workflow of C2C12 myotube differentiation, high insulin treatment and serum starvation. Differentiated myotubes were cultured in control (0 nM insulin) or hyperinsulinemic (2 or 200 nM insulin) medium for 16 hours and were analyzed before and after serum starvation. **(A’)** The insulin concentration in the medium at the end of 16-hour high insulin treatment (n=3). **(B)** Representative western blot images of phospho-AKT (T308, S473), total AKT, phospho-ERK1/2, and total ERK1/2. **(C)** Total AKT and **(D)** total ERK abundance under high insulin treatments before starvation (BS) and after starvation (AS). **(E)** phospho-AKT (T308), **(F)** phospho-AKT (S473) and **(G)** phospho-ERK1/2 measurements before starvation (BS), after starvation (AS), and stimulated by 0.2 or 2 nM insulin for 10 min after serum starvation. (n=6-9; ^#^ effect of hyperinsulinemia, ^$^ effect of starvation, ^&^ effect of acute insulin, × interaction between two factors, Mixed Effect Model.)

### Experimental Animals

Animal protocols were approved by the University of British Columbia Animal Care Committee. Isoform specific insulin deficient mice (*Ins1*^+/+^;*Ins2*^-/-^ and *Ins1*^+/-^;*Ins2*^-/-^) were randomly assigned to be fed *ad libitum* either a high-fat diet (Research Diets D12492, 20% protein, 60% fat, 20% carbohydrate content, energy density 5.21Kcal/g, Brunswick, NJ, US) or low-fat diet (Research Diets D12450B, 20% protein 10% fat, 70% carbohydrate content, energy density 3.82Kcal/g, Brunswick, NJ, US) for 4 weeks starting from 8 weeks old. Blood fasting glucose was measured using OneTouch Ultra2 glucose meters (LifeScan Canada Ltd, BC, Canada), and serum fasting insulin was assessed using mouse insulin ELISA kit (Alpco Diagnostics, Salem, NH, USA; detection limit 0.0324 – 1.19 nM), following 4-hour fasting. Lysates of gastrocnemius muscle from mice at 12 weeks old were used in this study.

### Western blot analyses

C2C12 myotubes or mice skeletal muscle (gastrocnemius) tissues were sonicated in RIPA buffer (50 mM β-glycerol phosphate, 10 mM HEPES, 1% Triton X-100, 70 mM NaCl, 2 mM EGTA, 1 mM Na_3_VO_4_, and 1 mM NaF) supplemented with complete mini protease inhibitor cocktail (Roche, Laval, QC), and lysates were resolved by SDS-PAGE. Proteins were then transferred to PVDF membranes (BioRad, CA) and probed with antibodies against p-ERK1/2 (Thr202/Tyr204) (1:1000, Cat. #4370), ERK1/2 (1:1000, Cat. #4695), p-AKT (Ser473) (1:1000, Cat. #9271), p-AKT (Thr308) (1:1000, Cat. #9275), AKT (1:1000, Cat. #9272), INSR-β subunit (1:1000, Cat. #3020S), p-INSRβ (Tyr1150/1151) (1:1000, Cat. #3024), FOXO1 (1:1000, Cat. #2880), p-FOXO1 (Thr24) (1:1000, Cat. #9464), all from Cell Signalling (CST), and β-tubulin (1:2000, Cat. #T0198, Sigma). The signals were detected by secondary HRP-conjugated antibodies (Anti-mouse, Cat. #7076; Anti-rabbit, Cat. #7074; CST) and Pierce ECL Western Blotting Substrate (Thermo Fisher Scientific). Protein band intensities were quantified with Image Studio Lite software (LI-COR).

### Surface Protein Biotinylation Assay

Biotinylation of surface proteins was performed as previously described (24) with modifications (Fig. 4A). In brief, cells were incubated with cell-impermeable EZ-Link-NHS-SS-biotin (300 μg/ml in PBS; Pierce) at 37°C for 2 min. Cells were then immediately placed on ice and washed with ice-cold 50 mM Tris-buffered saline (TBS) to remove excess biotin. For isolating surface proteins, cells were washed using ice-cold PBS and lysed in complete RIPA buffer (supplemented with cOmplete mini protease inhibitor cocktail (Roche, Laval, QC) and Na_3_VO_4_). For detecting internalized proteins, cells were washed with PBS and incubated in the serum-free medium supplemented with 0.2, 2, or 20 nM insulin at 37°C to stimulate INSR internalization. After certain time periods, cells were placed on ice, washed with ice-cold PBS, incubated with Glutathione solution (50 mM glutathione, 75 mM NaCl, 1 mM EDTA, 1% BSA, 75 mM NaOH) for 20 min to strip the remaining surface biotin, washed with excess PBS, and lysed in complete RIPA buffer. Lysates were quantitated and incubated with NeutrAvidin beads (Pierce) overnight at 4 °C to isolate biotinylated surface or internalized proteins. Biotinylated proteins were eluted from the NeutrAvidin beads by boiling in Blue Loading Buffer (CST) containing 50 mM DTT for 5 min. Surface or internalized INSR in eluent and total INSR in lysates were detected in Western blot analysis.

### siRNA knockdown in C2C12 myoblasts

All siRNAs are from Thermo Fisher Scientific with the specific Assay IDs as follows: *Foxo1* (MSS226201), *Sin3a* (151684), *Elf1* (157302), *Mxi1* (68202), *Myc* (68302), *Ets1* (101877), *Hcfc1* (158001), *Nrf1* (68266), *Jund* (67635), *Ctcf* (60925), *Max* (155266), *Maz* (501159), Silencer Cy3-labeled Negative Control No.1 siRNA (AM4621). siRNAs were transfected into C2C12 myoblasts using the Lipofectamine RNAiMAX reagent (Invitrogen) according to the manufacturer’s instructions with 50 pmol of siRNA and 4 µl of transfection reagent per well of a 12-well plate.

### RNA isolation and quantitative real-time PCR analysis

Total RNA was isolated from both control and high insulin-treated C2C12 myotubes before and after serum starvation or C2C12 myoblasts post siRNA transfection using the RNEasy mini kit (Qiagen). cDNA was generated by reverse transcription using qScript cDNA synthesis kit (Quanta Biosciences, Gaithersburg, MD, USA). Transcript levels of target genes in the equal amount of total cDNA were quantified with SYBR green chemistry (Quanta Biosciences) on a StepOnePlus Real-time PCR System (Applied Biosystems). All data were normalized to *Hprt* by the 2^−^Δ^Ct^ method. The following primers are used in qPCR: *Insr-A*/*B* forward 5’-TCCTGAAGGAGCTGGAGGAGT-3’, *Insr-A* reverse 5’-CTTTCGGGATGGCCTGG-3’, *Insr-B* reverse 5’-TTCGGGATGGCCTACTGTC-3’ (25); *Insr* (in siRNA experiments) forward 5’-TTTGTCATGGATGGAGGCTA-3’ and reverse 5’-CCTCATCTTGGGGTTGAACT-3’ (26) *Igf1r* forward 5’-GGCACAACTACTGCTCCAAAGAC-3’ and reverse 5’-CTT TATCACCACCACACACTTCTG-3’ (25); *Hprt* forward 5’-TCAGTCAACGGGGGACATAAA-3’ and reverse 5’-GGGGCTGTACTGCTTAACCAG -3’ (27); *Foxo1* forward 5’-CCCAGGCCGGAGTTTAACC-3’ and reverse 5’-GTTGCTCATAAA GTCGGTGCT-3’, *Tbp* forward 5’-AGAACAATCCAGACTAGCAGCA-3’ and reverse 5’-GGGAACTTCACATCACAGCTC-3’, *Nrf1* forward 5’-TATGGCGGAAGTAATGAAAGACG-3’ and reverse 5’-CAACGTAAGCTCTGCCTTGTT-3’, *Jund* forward 5’-GAAACGCCCTTCTATGGCGA -3’ and reverse 5’-CAGCGCGTCTTTCTTCAGC-3’, *Ctcf* forward 5’-GATCCTACCCTTCTCCAGATGAA-3’ and reverse 5’-GTACCGTCACAGGAACAGGT-3’, *Mxi1* forward 5’-AACATGGCTACGCCTCATCG-3’ and reverse 5’-CGGTTCTTTTCCAACTCATTGTG-3’, *Elf1* forward 5’-TGTCCAACAGAACGACCTAGT-3’ and reverse 5’-CACACAAGCTAGACCAGCATAA-3’, *Ets1* forward 5’-TCCTATCAGCTCGGAAGAACTC-3’ and reverse 5’-TCTTGCTTGATGGCAAAGTAGTC-3’, *Maz* forward 5’-GCCCCAGTTGCATCTGTCTT-3’ and reverse 5’-CTTCGGAGGTTGTAGCCGTT-3’, *Max* forward 5’-ACCATAATGCACTGGAACGAAA-3’ and reverse 5’-GTCCCGCAAACTGTGAAAGC-3’, *Myc* forward 5’-ATGCCCCTCAACGTGAACTTC-3’ and reverse 5’-CGCAACATAGGATGGAGAGCA-3’, *Hcfc1* forward 5’-CGGCAACGAGGGGATAGTG-3’ and reverse 5’-TAGGCGAGTACCATCACACAC-3’, *Sin3a* forward 5’-GCCTGTGGAGTTTAATCATGCC-3’ and reverse 5’-CCTCTTGCTCAGTCAAAGCTG-3’, *Irs2* forward 5’-CTGCGTCCTCTCCCAAAGTG-3’ and reverse 5’-GGGGTCATGGGCATGTAGC-3’ (PrimerBank, https://pga.mgh.harvard.edu/primerbank/index.Html).

### RNA sequencing and bioinformatic analysis

Total RNA isolated from both control and 200 nM insulin-treated C2C12 myotubes before and after serum starvation (4 groups, n = 5 each group) were sequenced by BRC Sequencing Core at the University of British Columbia. Sample quality control was performed using the Agilent 2100 Bioanalyzer. Qualifying samples were then prepped following the standard protocol for the NEBNext Ultra II Stranded mRNA (New England Biolabs). Sequencing was performed on the Illumina NextSeq 500 with Paired-End 42bp × 42bp reads. Sequencing data were demultiplexed using Illumina’s bcl2fastq2. De-multiplexed read sequences were then aligned to the Mus Musculus mm10 reference sequence using STAR aligner (28). Assembly was estimated using Cufflinks (http://cole-trapnell-lab.github.io/cufflinks/) using methods available on the Illumina Sequence Hub and with default settings.

Raw counts of the gene reads were filtered for minimal expression by only keeping genes with more than 5 raw reads in more than 5 samples. After normalizing counts by variance stabilizing transformation, differential expression analysis was performed using DESeq2 package (29). Raw counts of the mouse skeletal muscle data sets (IRMOE (30) and Clamp (22)) were analyzed using the same criteria. Kyoto Encyclopedia of Genes and Genomes (KEGG) pathway enrichment analyses were performed using the R package clusterProfiler (31). Reactome pathway enrichment analyses were performed using the R package ReactomePA (32). The background gene list was set to be all the detectable genes passing the minimal expression filter. Transcription factor (TF)-gene network was derived from the ENCODE ChIP-seq data and illustrated by a visual analytic platform NetworkAnalyst 3.0 (http://www.networkanalyst.ca/) (33). Top 30 TFs with the highest degree of connections (rank order) with differentially expressed genes were selected for siRNA knockdown. Redundant pathways, containing many overlapping genes, were omitted in the figures but are included in supplementary tables.

### Human muscle transcriptomic meta-analyses

We carried out a meta-analysis using three independent human skeletal muscle transcriptomic data sets (n=488) (Table 1). In all cases, insulin values were log transformed prior to any analysis and were normally distributed under these conditions. Pearson correlation coefficient (R) between fasting insulin and normalized gene expression was calculated. Genes with significant correlation (p<0.05, | R| >0.2) were selected for further comparisons.

**Table 1.** Phenotypes of human cohorts.

| Data sets | Insulin (pM) |  | Glucose (mM) |  | Age |  | Sex |  |
| --- | --- | --- | --- | --- | --- | --- | --- | --- |
|  | Mean | SD | Mean | SD | Mean | SD | Female | Male |
| SMP | 68.1 | 42.7 | 5.02 | 0.69 | 43.3 | 15.0 | 102 | 89 |
| FUSION | 59.1 | 34.9 | 6.27 | 0.78 | 60.0 | 7.6 | 115 | 163 |
| Møller *et al* | 176.8 | 150.7 | 6.99 | 1.41 | 60.8 | 6.6 | 7 | 12 |

We utilised a large tiling-type microarray data set, referred to as SMP (<u>S</u>TRRIDE-PD and <u>M</u>ETA<u>P</u>REDICT studies), which consisted of 191 sedentary individuals with increased body mass index (BMI) and/or impaired glucose tolerance. Gene expression profiles (HTA 2.0 array) and plasma insulin values (K6219 Dako high-sensitivity enzyme linked immunosorbent assay (ELISA)) were generated in a single core lab (34). Gene expression was normalized using IRON (35) producing log2 transformed data for 53,032 probe-sets representing the detectable protein coding Ensembl transcripts.

Two RNA-seq muscle tissue cohorts were utilized. A large study, FUSION (Finland-United States Investigation of NIDDM Genetics Study; dbGaP accession phs001048.v2.p1), consisted of individuals (N=299) classified into normal glucose tolerance (NGT), impaired fasting glucose (IFG), impaired glucose tolerance (IGT), or type 2 diabetes (T2D). Two hundred and seventy eight RNAseq samples (HiSeq2000 using 101bp paired-end reads) passed QC and were correlated with fasting insulin with muscle RNA. Insulin was measured in serum using the Architect chemiluminescent microparticle immunoassay. A second, case-control human RNA-seq data set (Møller *et al*) is a small cohort (N=19) consists of age-matched health subjects (Control), T2D subjects with severe insulin resistance under insulin injection (T2D-SI group), or oral anti-diabetic drug (T2D-OAD group)(36, 37). Serum insulin was analyzed using time-resolved immunofluorometric assay (AutoDELFIA, PerkinElmer, Finland). Raw counts of the RNA-seq gene reads were normalized by variance stabilizing transformation.

### Statistics

Data were presented as mean ± SEM in addition to the individual data points. All data were analyzed using R Studio 3.4.1. A significance level of adjusted p < 0.05 was used throughout. All western blot quantifications (protein band intensity) were analyzed using linear regression modeling (38). Linear mixed effect models (R package – lme4) were fitted using restricted maximum likelihood (38, 39). Predictor variables were included as fixed effects and sample IDs were included as random effects. Mixed effect modeling was used to account for repeated sample measurements and missing data (38). Where the random effect was not significant, linear fixed effect modeling was used. Heteroscedasticity and normality of residuals were analyzed used Levene’s test and the Shapiro–Wilk test, respectively. Predictor variables, insulin treatment (overnight and acute) and time, were treated as ordinal factors and continuous factors, respectively. The outcome variable, protein band intensity, was treated as a continuous factor and log-transformed when residuals are not homoscedastic and/or normally distributed. Multiple comparison p-values were adjusted using the Tukey method.

## Data and Code Availability

The new RNA-seq data from the cell experiments are available under GEO accession GSE147422. All remaining data are contained within the article and can be shared upon request or at GEO (GSE154846). Code for all the analyses is available at GitHub deposit https://github.com/hcen/Hyperinsulinemia_muscle_INSR.

## Results

### Hyperinsulinemia induces insulin resistance in muscle cells *in vitro*

Circulating insulin in humans oscillates in a range between approximately 0.01 nM and 2 nM, and fasting insulin less than 0.085 nM is considered normal (34, 40, 41). To establish an *in vitro* modeling of hyperinsulinemia condition, which is referred to as hyperinsulinemia, we incubated differentiated mouse C2C12 myotubes for 16 hours in a physiologically high insulin dose of 2 nM or supraphysiological high dose of 200 nM (Fig. 1A). Hyperinsulinemia was confirmed after treatment with high insulin (Fig. 1A’). After 6 hours of serum starvation, insulin signaling was assessed by measuring the phosphorylation of AKT and ERK proteins, two major insulin signaling nodes (42). Since alterations in the basal state of the insulin signal transduction network have also been reported in hyperinsulinemic humans and animals (43), we also measured the effects of hyperinsulinemia on AKT and ERK phosphorylation before and after serum starvation (BS and AS). These experiments showed that total AKT protein was downregulated by prolonged 200 nM, but not 2 nM, insulin treatment, while ERK abundance was not significantly changed (Fig. 1B-D). After prolonged 200 nM insulin exposure - and before serum starvation - AKT phosphorylation at threonine (T) 308 and serine (S) 473 was significantly elevated, while ERK phosphorylation was unaffected (Fig. 1E-G). Of note, phosphorylation of ERK1/2 was increased by serum starvation alone, as previously reported in other cell types (Fig. 1G) (44). Acute AKT and ERK signaling in the context of 2 nM acute insulin was significantly reduced by hyperinsulinemia treatment in an insulin dose-dependent manner (Fig. 1E-G). We also characterized the insulin dose- and time-dependent signaling in our *in vitro* hyperinsulinemia model (200 nM insulin). Phosphorylation of AKT and ERK1/2 were significantly reduced under 0.2, 2, or 20 nM insulin stimulations (Fig. S1A,B). Together, these results establish a robust muscle cell insulin resistance response induced by 16 hours of hyperinsulinemia.

### Insulin signaling genes are modulated by hyperinsulinemia and serum starvation in muscle cells

To further investigate the molecular mechanisms of hyperinsulinemia-induced insulin resistance, we conducted RNA sequencing (RNA-seq) on cells exposed to prolonged insulin and serum starvation. We compared the transcriptomes across 4 treatment groups (n=5 per group, 0 nM or 200 nM insulin, both before and after serum starvation). Principal component analysis (PCA) of global gene expression showed that the majority of experimental variation was related to hyperinsulinemia (PCA1), while the impact of starvation was evident in PCA2 (greatly enhanced in the hyperinsulinemia conditions) (Fig. 2A). There were 2882 up-regulated and 2506 down-regulated genes before starvation (BS, 200 nM vs 0 nM) (Table S1), and the top 50 most significantly altered genes were shown in Figure S2A. We analyzed their functions using Reactome and KEGG pathway enrichment analyses (Fig. 2B,C, Fig. S2B, Table S2,3). The genes upregulated by hyperinsulinemia (BS, 200 vs 0 nM) enriched pathways related to cell cycle, RNA biology, translation, and glucose metabolism (Fig. 2B, Table S2,3), while the downregulated genes enriched various signaling pathways (Fig. 2C,S2B, Table S2,3).

**Figure 2.**
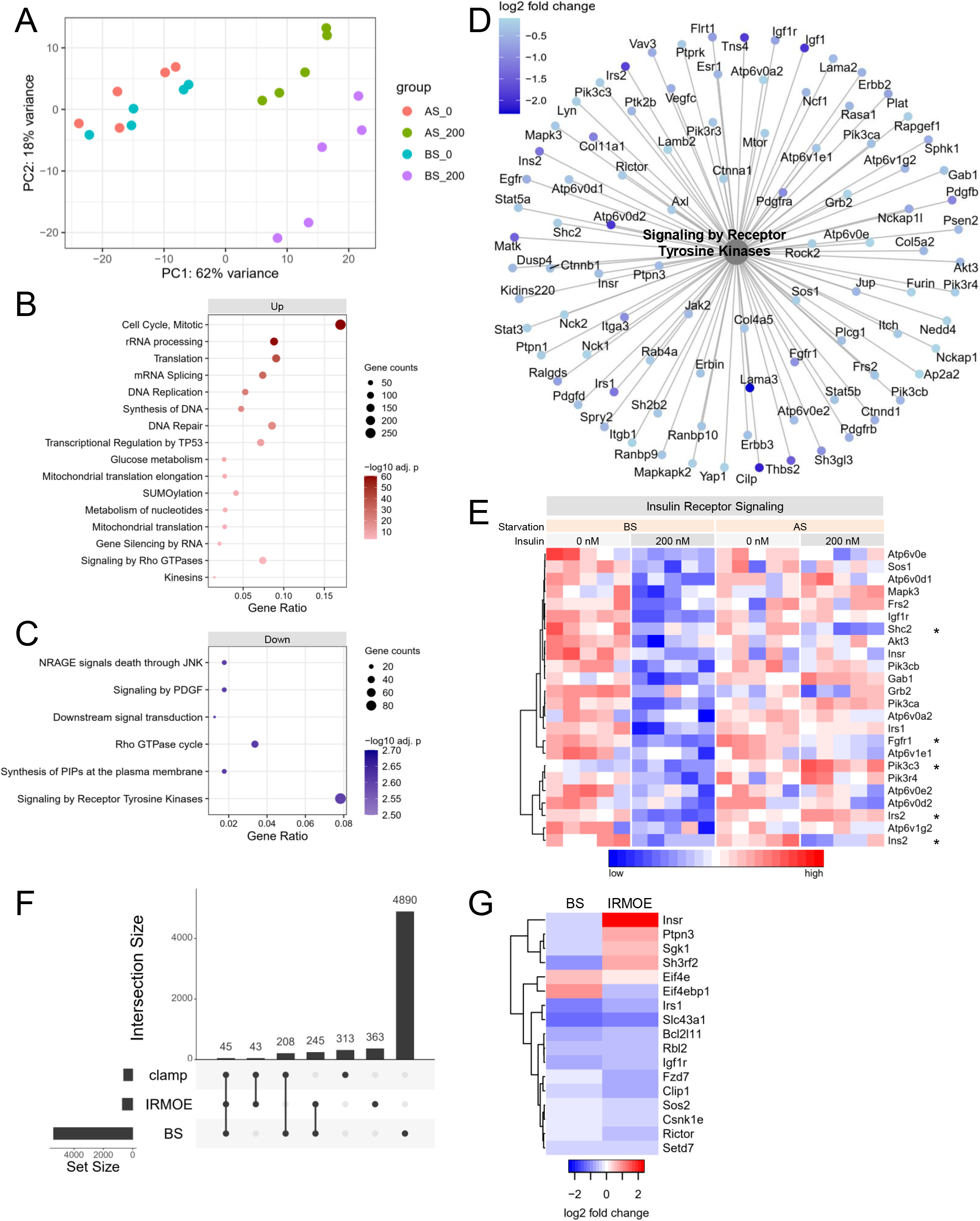
Transcriptomic analysis of hyperinsulinemia and serum starvation. **(A)** Principle component analysis (PCA) of RNAseq data from 4 groups of treatments, including 0 nM or 200 nM prolonged insulin both before and after serum starvation. Selected top Reactome pathways enriched from **(B)** upregulated or **(C)** downregulated genes by hyperinsulinemia before starvation (BS, 0 vs 200 nM insulin). **(D)** Differentially expressed genes (BS, 200 vs 0 nM) enriched under Reactome pathway “Signaling by Receptor Tyrosine Kinases”. **(E)** The normalized gene expressions of downregulated insulin signaling genes before starvation (BS, 200 vs 0 nM). * Genes that are also differentially expressed after starvation (AS, 200 vs 0 nM). **(F)** The number of DE genes shared between or *in vitro* study (BS), IRMOE study, and hyperinsulinemic clamp study (clamp). **(H)** The log2 fold change of the genes of interest that consistently altered in our *in vitro* model (BS, 200 vs 0 nM) and IRMOE muscle.

Starvation (200 nM, BS vs AS) changed some of these pathways in the opposite direction, while some pathways remained to be downregulated after starvation (AS, 200 vs 0 nM) (Table S2). For example, many upregulated genes related to glucose metabolism were upregulated by hyperinsulinemia, such as *G6pc3, Hk1, Pck2* and *Aldoa*, some of which were recovered by starvation (Fig. S2C). Many interesting downregulated genes were linked to Reactome pathway “Signaling by Receptor Tyrosine Kinase” (FDR=0.002) (Fig. 2C), and a subset of genes of interest related to insulin receptor signaling were highlighted (Fig. 2D). Most of these insulin signaling genes were recovered after starvation, such as *Insr* and *Irs1*; some remain downregulated, such as *Shc2* and *Fgf1r*; some were upregulated instead, such as *Irs2* and *Pik3c3* (Fig. 2D). Despite the recovery effect of starvation, a large proportion of differentially expressed genes before starvation remained to be altered in the same direction after starvation (Fig. S2D), demonstrating the long-lasting effects of hyperinsulinemia. Overall, prolonged hyperinsulinemia and insulin removal revealed strong, reciprocal transcriptomic effects. Many insulin signaling genes were reprogramed by hyperinsulinemia, which may contribute to the insulin resistance in our model.

To support the relevance of our hyperinsulinemia-induced transcriptomic changes to *in vivo*, we compared the RNA responses in our *in vitro* model with two mouse skeletal muscle systems with sustained insulin signaling. One recent study with insulin receptor muscle over-expression (IRMOE) demonstrated similarities with our *in vitro* model, such as increased basal INSR and AKT phosphorylation and impaired insulin-stimulated AKT phosphorylation – indicative of a significant level of post-receptor insulin resistance (30). The second study used the hyperinsulinemic-euglycemic clamp, in which ∼1 nM insulin plasma insulin was maintained for 3 hours (high insulin vs saline) (22), which represents relatively acute hyperinsulinemia. Compared with IRMOE and clamp studies, our *in vitro* hyperinsulinemia model had different subsets of overlapping differentially expressed genes (Fig.2G, S3A,B, Table S4). Among the common differentially expressed genes between our *in vitro* model and the IRMOE system, several genes of interest were consistently altered, such as *Irs1, Igf1r, Sos*, and *Rictor* (Fig. 2G). It appears that our *in vitro* model represents many features observed with excess insulin signaling *in vivo*.

### Transcriptomic analysis of human skeletal muscle reveals genes associated with fasting insulin including *INSR*

We established above that a key feature of excess insulin signaling was the down-regulation of components of proximal insulin signaling, including the insulin receptor gene (*Insr*). To explore if these specific molecular responses are consistent with human clinical responses, we modeled the relationship between fasting insulin and the gene expression in human skeletal muscle (Table 1) (34, 36, 45). These human cohorts represent the full range of insulin resistance and cover normal glucose tolerance (34, 36), pre-diabetes and diabetes (34, 36) to extreme obesity-related insulin resistance (45). One of our human cohorts had a higher range of fasting insulin (Fig. S3C) because the study included diabetic subjects with severe insulin resistance needing insulin injection (T2D-SI group) (36). Although correlation analysis is often not appropriate for small case-controlled studies, the range of insulin values overlapped and we observed that *INSR* and *IRS2* gene expression levels had a negative trend with fasting insulin (Fig. 3A). Using the other two large human cohorts, after cross-referencing with orthologous mouse genes in our hyperinsulinemia model, we identified genes that were significantly correlated with fasting insulin and differentially expressed in our hyperinsulinemia model (Fig. 3B,C, Table S5). Compellingly, *INSR* was negatively correlated with fasting insulin in all data sets (Fig. 3C), consistent with our *in vitro* model and previous reports using a HOMA2-IR model (34). *INSR* and insulin also had a negative correlation in the normal glucose tolerance group in the FUSION human cohort (Fig. 3D), suggesting that the association is independent of pathological insulin resistance or diabetes.

**Figure 3.**
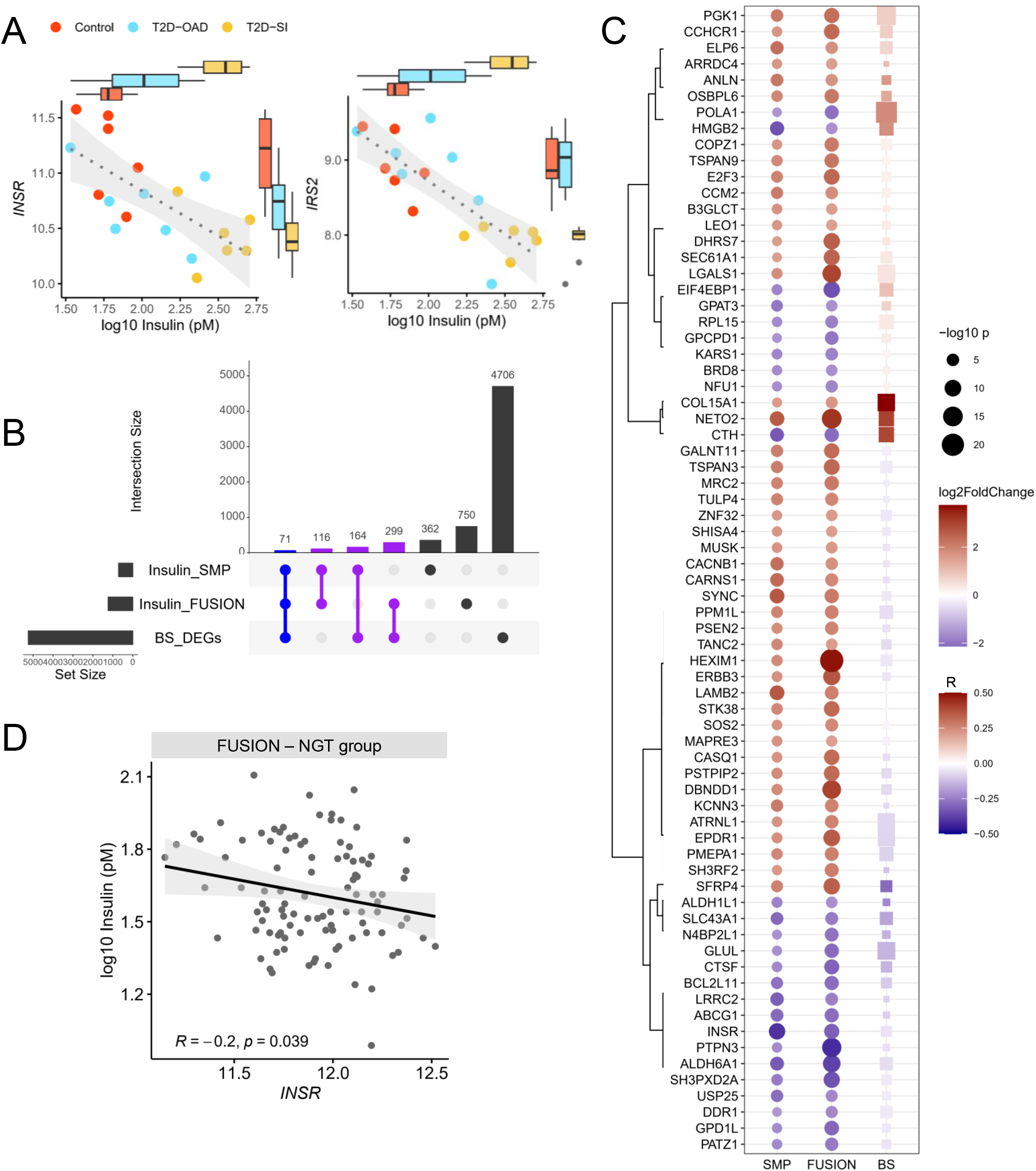
Correlation between fasting insulin and human transcriptome highlights insulin receptor expression. **(A)** The INSR and IRS2 expressions and the corresponding fasting insulin levels in control subjects, diabetic subjects with oral antidiabetic drug (T2D-OAD) or with severe insulin resistance (T2D-SI). **(B)** The number of genes correlated with fasting insulin or differentially expressed in our *in vitro* model (BS_DEGs) shared between data sets. Genes in blue intersection and their Pearson correlation coefficient (R) in human data sets or log2 fold change in our in vitro hyperinsulinemia model (BS) are displayed in **(C). (D)** The correlation between INSR and insulin in normal glucose tolerance (NGT) group in the FUSION human cohort.

### Hyperinsulinemia reduces both *Insr* isoform A and B alongside FOXO1 inhibition

Loss of *INSR* expression reflects that it is a critical component as the very start of insulin signaling. Downregulation of in *Insr* expression as well as alternative splicing in some (46-48), but not all analysis (34) has been associated with hyperinsulinemia. Therefore, we carried out a more detailed analysis of transcription from the *Insr* gene in our model, using qPCR. Both *Insr-A* and *Insr-B* mRNA were equally and robustly downregulated after hyperinsulinemia and recovered by serum starvation (Fig. 4A). The ratio of *Insr-A* and *Insr-B* mRNA was not affected by hyperinsulinemia or serum starvation in cultured muscle cells (Fig. 4B). Insulin-like growth factor 1 receptor (*Igfr1*), which has a similar structure and signaling mechanism as INSR, was also reduced by hyperinsulinemia at the transcriptional level (Fig. 4C). Therefore, the loss of *Insr* was not compensated by *Igf1r* under these conditions nor related to differential splicing of the INSR gene, consistent with *in vivo* observations using the largest clinical cohorts (34).

Forkhead box protein O1 (FOXO1) is a known transcriptional regulator of the *Insr* gene and is also a key mediator of insulin signaling (49-51). In *Drosophila* and mouse myoblasts, FOXO1 activity is necessary and sufficient to increase *Insr* transcription under serum fasting and reverse this effect in the presence of insulin (50). We noted that genes downregulated by hyperinsulinemia belonged to the “FOXO signaling pathway” (FDR=9.95×10^-4^), and most of these genes were recovered by starvation (Fig. 4D. S2B). Therefore, we sought to determine the activity of FOXO1 in our hyperinsulinemic model (Fig. 4E). Insulin increased FOXO1 phosphorylation on T24, which is an AKT-associated event known to exclude FOXO1 from the nucleus, decreasing its transcriptional activity (52), without altering total FOXO1 abundance (Fig. 4E). T24 phosphorylation of FOXO1 decreased after starvation (Fig. 4E), consistent with our observed effects on AKT phosphorylation and *Insr* transcription. Our data, therefore, support the work of other groups indicating a role for FOXO1 in *Insr* gene expression. However, knocking down *Foxo1* mRNA by 40% did not alter *Insr* mRNA level in C2C12 myoblast (n=5, Fig. 4F), indicating redundancy in terms of transcriptional control of *Insr*, or that the remaining 60% of *Foxo1* expression was sufficient to maintain *Insr* gene expression.

**Figure 4.**
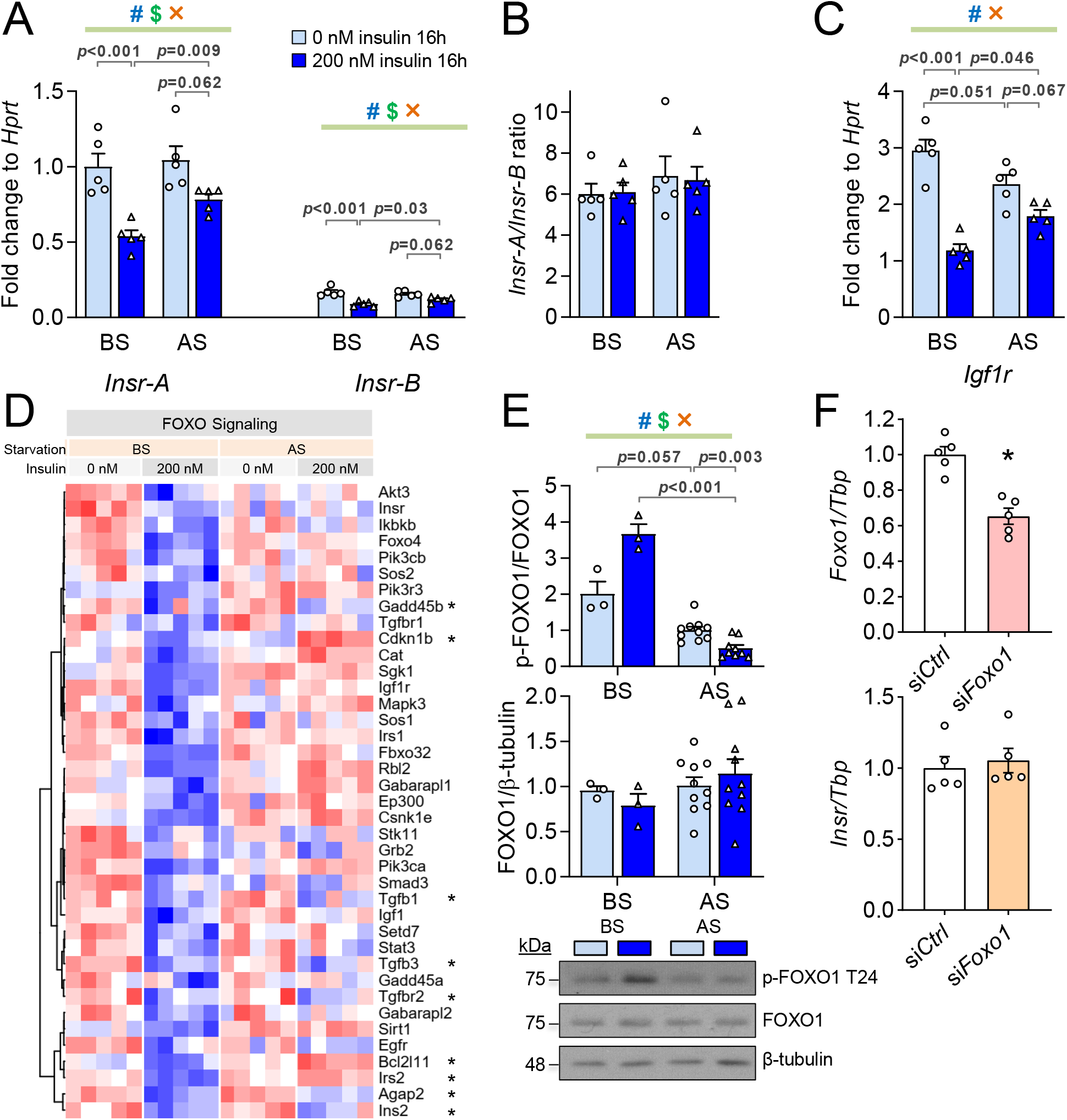
Effects of prolonged hyperinsulinemia and starvation on *Insr* transcription and FOXO1 phosphorylation *in vitro*. **(A)** The mRNA levels of *Insr* isoform A or B *(Insr-A or B)* before and after starvation (BS and AS) assessed by qPCR. **(B)** *Igf1r* mRNA level. **(C)** The ratio of *Insr-A* to *Insr-B* mRNA (n=5). (D) Normalized counts of downregulated genes under KEGG pathway “FOXO signaling” before starvation (BS, 200 vs 0 nM). * Genes that are also differentially expressed after starvation (AS, 200 vs 0 nM). **(E)** Total and T24 phosphorylation of FOXO1 (n = 3 in BS group, n = 10 in AS group). (F). **(C)** mRNA levels of *Foxo1* and *Insr* after knocking down Foxo1 by siRNA (n=5, * *p*<0.05). (^#^ effect of hyperinsulinemia, ^$^ effect of starvation, × interaction between two factors, Mixed Effect Model.)

### Hyperinsulinemia reduces INSR protein abundance but not its phosphorylation or internalization

To further examine the direct effects of hyperinsulinemia on the proximal stages of insulin signaling, we examined INSR abundance, phosphorylation and internalization in cultured muscle cells. Consistent with the reduction in *Insr* mRNA, total INSR protein abundance was robustly decreased in both 2 and 200 nM hyperinsulinemia groups in an insulin dose-dependent manner (Fig. 5A, B). Serum starvation slightly recovered the INSR downregulation in 200 nM insulin group (Fig. 5B). These results clearly demonstrated that prolonged insulin directly modulates INSR abundance in this cell system – consistent with *in vivo* clinical correlations.

**Figure 5.**
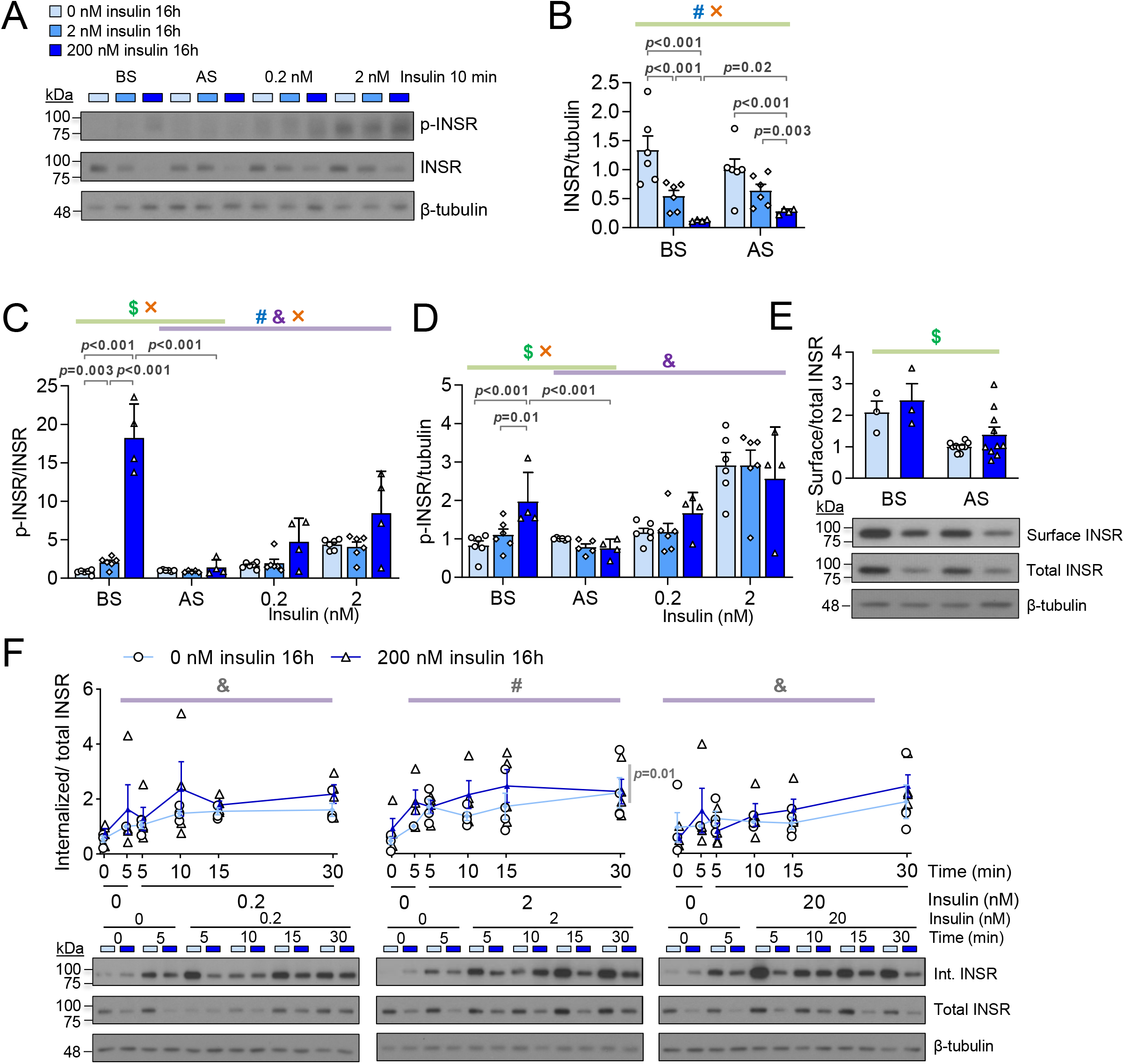
Effects of hyperinsulinemia and serum starvation on INSR abundance, phosphorylation and internalization. **(A)** Representative western blot images of phospho-INSR (Y1150/1151) and total INSR. **(B)** The level of total INSR protein before or after serum starvation (n=4-6). The ratio of phospho-INSR (Y1150/1151) to total INSR **(C)** or tubulin **(D)** before starvation (BS), after starvation (AS), and stimulated by 0.2 or 2 nM insulin for 10 min after serum starvation (n=4-6). **(E)** The ratio of surface to total INSR ((n = 3 in BS group, n = 10 in AS group). **(F)** The ratio of internalized to total INSR over 30 min under 0.2, 2 or 20 nM acute insulin stimulation (n = 4). (^#^ effect of hyperinsulinemia, ^$^ effect of starvation, × interaction between two factors, Mixed Effect Model.)

We also examined INSR tyrosine 1150/1151 autophosphorylation, which is an early step of insulin signaling that recruits IRS and SHC, leading to PI3K-AKT or RAS-ERK activation (42). Before starvation, both hyperinsulinemia groups had increased INSR phosphorylation, suggesting that there was continuous insulin signaling during the high insulin treatments (Fig. 5C,D). Serum starvation completely reversed INSR hyperphosphorylation (Fig. 5C,D). While INSR phosphorylation was not significantly different after 10 min of acute insulin stimulation (Fig. 5C,D), analysis of dose- and time-dependent insulin signaling revealed a tendency for increased phosphorylated-to-total INSR ratio in insulin-stimulated cells exposed overnight to 200 nM insulin (Fig. S1C). The increased INSR phosphorylation per receptor was offset by the reduced INSR number, leading to a decreased phospho-INSR-to-tubulin ratio (i.e. the overall INSR phosphorylation events per cell) (Fig. S1C). These data indicate that the were not defects in INSR phosphorylation upon acute insulin stimulation in our system, but the INSR abundance could limit the overall INSR phosphorylation.

Impaired INSR endocytosis has been implicated in insulin resistance (53, 54). Many genes related to endocytosis pathways were downregulated by hyperinsulinemia (BS, 200 vs 0 nM) and recovered by starvation (200 nM, AS vs BS) (Fig. S4A). Therefore, basal surface INSR, as well as dose- and time-dependent INSR internalization were examined in our hyperinsulinemia model using a surface biotinylation assay (Fig. S4B). Serum starvation slightly decreased the surface-to-total INSR ratio, while hyperinsulinemia had no significant effects (Fig. 4E). Upon acute insulin stimulation, the internalized INSR to total INSR ratio did not have evident differences except for a small increase when stimulated by 2 nM insulin (Fig. 4F). Therefore, hyperinsulinemia-induced insulin resistance may be mediated by a reduction in total INSR that results in a proportional reduction in INSR protein at the cell surface. The fraction of INSR internalized during acute insulin signaling seemed to be recalibrated instead of drastically affected by hyperinsulinemia under these conditions. Collectively, our experiments suggest that hyperinsulinemia-induced insulin resistance in muscle cells is mediated by a reduction in total INSR, and not primarily by affecting its activity or internalization.

### Circulating insulin negatively correlates with INSR protein level in vivo

To further extend our *in vitro* studies, we examined the relationship between *in vivo* insulin concentration and muscle INSR protein abundance in mice. As in our previous studies (13), insulin gene dosage was manipulated to generate variance in circulating insulin. The mice were fed with a high-fat diet (HFD) known to induced pronounced hyperinsulinemia (13, 55) or a low-fat diet (LFD) for 12 weeks. Fasting insulin and glucose were higher in male mice. These experiments showed that, in male mice, INSR protein abundance negatively correlated with both fasting insulin and fasting glucose in the HFD group (Fig. 6A,B). However, the LFD group had a negative correlation between INSR level and insulin, with no correlation between INSR and glucose (Fig. 6A,B). On the other hand, in female mice, fasting insulin levels were near or at the lower detection limit, reducing measurement dynamic range and contributing to the lack of correlation with INSR (Fig. 6A). Fasting glucose levels did not correlate with INSR (Fig. 6B). Together, these data support the concept that insulin, independent from glucose, negatively regulates INSR levels in skeletal muscle. This is consistent with our *in vitro* hyperinsulinemia model and our previous *in vivo* data demonstrating improved insulin sensitivity over time in mice with genetically reduced insulin production (12). These data also suggest an interaction between insulin, glucose and INSR that is dependent on the conditions of the HFD. The source of the aforementioned sex differences may again reflect insulin dose-dependent effects and requires further investigation.

**Figure 6.**
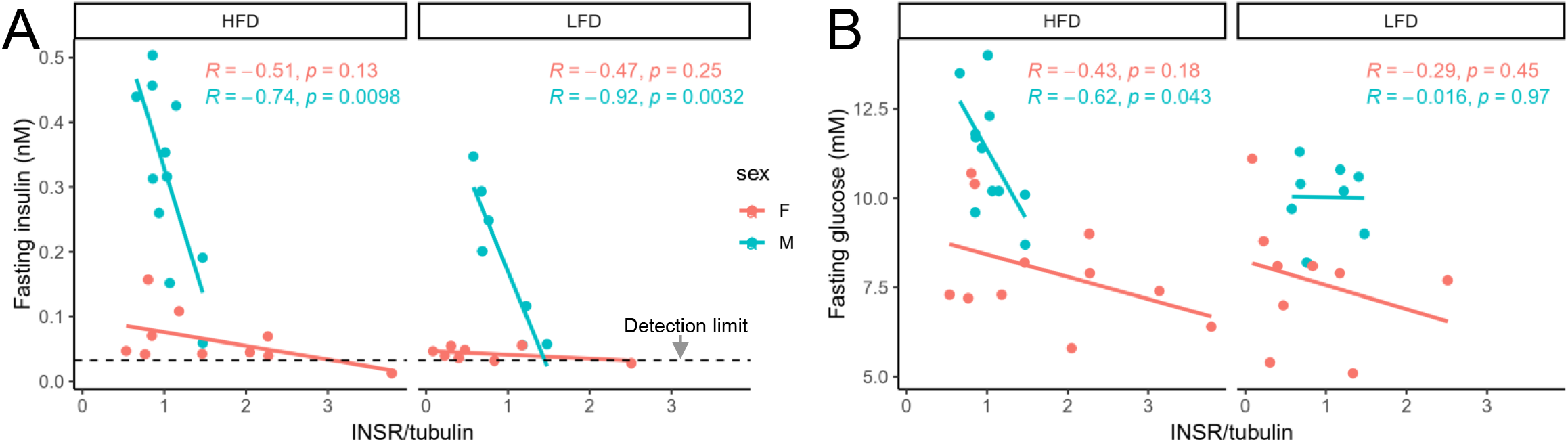
*In vivo* correlation between INSR abundance fasting insulin and glucose in skeletal muscle. **(A)** Pearson correlation between fasting insulin and INSR protein abundance in female (F) or male (M) HFD or LFD-fed mice. **(B)** Pearson correlation between fasting glucose and INSR protein abundance in female (F) or male (M) HFD or LFD-fed mice. (n = 7-11)

**Figure 7.**
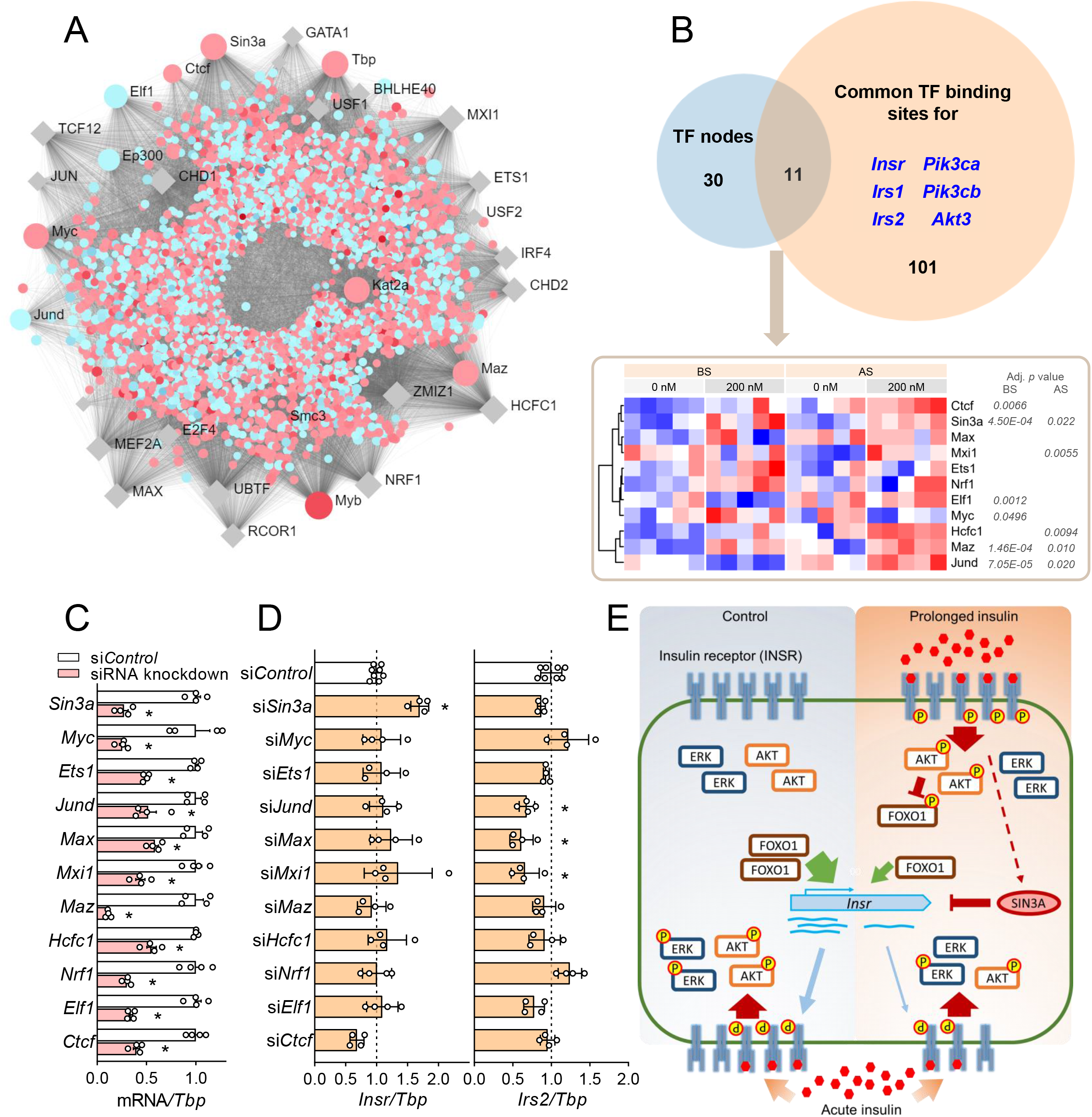
Identification of upstream transcription factors mediating the overall transcriptomic changes and insulin receptor expression. **(A)** TF-gene network predicting upstream transcriptional regulators of differentially expressed genes by hyperinsulinemia (BS, 200 vs 0 nM). Names of the top 30 TFs are labeled. Genes that are up or down-regulated are shown in red or blue dots, respectively. TFs that are not differentially expressed are in grey rhombus. **(B)** Common TFs between the top 30 TF nodes from (A) and the TF binding sites for several key genes in insulin signaling pathways that are downregulated by hyperinsulinemia. Adjusted p values of differentially expressed TFs (200 vs 0 nM, both BS and AS) are labeled next to the heatmap. **(C)** mRNA levels of the common TFs in (B) after siRNA knockdown. **(D)** The effects of each siRNA knockdown on *Insr* and *Irs2* mRNA levels. (n=4. * *p*<0.05, 1-ANOVA followed by Dunnett multiple comparison test against si*Control*.) **(E)** Graphic summary of our current model. Hyperinsulinemia induced sustained phosphorylation of INSR and AKT, which resulted in the inhibition of FOXO1 leading to reduced *Insr* transcription. Downregulated INSR and post-receptor components resulted in reduced insulin signaling upon acute insulin stimulation. SIN3A, which was upregulated by hyperinsulinemia, might repress *Insr* transcription.

### Novel transcription factors regulate *Insr* expression and transcriptomic remodeling by insulin

We identified upstream transcriptional regulatory proteins by examining transcription binding sites in the differentially expressed genes using ENCODE ChIP-seq data (Fig. 8A). The top 30 transcriptional factors with high degrees of connections to altered genes (Fig. 8A), were compared to common transcription factor binding sites of several differentially expressed genes that encode key proteins in insulin signaling (Fig. 8B). The resulting 11 transcription factors all have binding sites near the transcription start sites of all the selected insulin signaling genes based on the ENCODE ChIP-seq data and were deemed candidates to affect the transcriptional changes in *Insr* and the other selected insulin signaling genes during hyperinsulinemia (Fig. 8B). Among them, *Sin3a, Myc* and *Ets1* were upregulated by hyperinsulinemia (Fig. 8B). To investigate the role of these transcription factors on *Insr* expression, we conducted siRNA knockdown for each transcription factor, with knockdown efficiencies varying between 45% and 85% (Fig. 8C). *Sin3a* knockdown (∼70%) resulted in a significant increase in *Insr* mRNA, indicating that this transcription factor has a repressive effect (Fig. 8D). In addition, we assessed the expression of *Irs2*, which had a larger fold change than *Insr* and was one of the most significantly altered genes upon hyperinsulinemia and starvation (Fig. S2A). Knockdown of *Jund, Max* and *Mxi1* downregulated *Irs2*, which suggested that these transcription factors are involved in *Irs2* transcription and may contribute to insulin resistance (Fig. 8D). In conclusion, we identified transcription factors for *Insr* and *Irs2* genes among the predicted upstream transcriptional regulators.

## Discussion

The goal of this study was to explore the mechanisms of hyperinsulinemia-induced insulin resistance in skeletal muscle cells with a focus on transcriptomic changes. We demonstrated that prolonged physiological and supraphysiological hyperinsulinemia induced a reduction of AKT and ERK signaling (Fig. 8E). Remarkably, while serum starvation partially reversed the effects of overnight hyperinsulinemia, much of the impaired acute insulin signaling and transcriptomic remodeling was sustained after 6 hours of insulin withdrawal and serum starvation, suggesting that stable molecular changes underlie these differences. The effects of prolonged hyperinsulinemia were insulin dose-dependent from the physiological to the supraphysiological range. We demonstrated that the impaired insulin response in our system can be partially accounted for by INSR downregulation at the transcription level and also used transcriptomic profiling to discover new factors that regulate insulin signaling in our system including SIN3A, JUND, MAX and MXI1. These experiments showcased many genes that were reprogramed by hyperinsulinemia and insulin removal.

Our *in vitro* cell culture model provided a robust and controlled system for examining the direct effects of excess insulin, and insulin withdrawal, on multiple components of insulin signaling in muscle cells. Our results are consistent with other *in vitro* cell culture systems designed to examine the effects of hyperinsulinemia. For example, reduced AKT and ERK signaling and INSR abundance were also reported in hyperinsulinemia-treated β-cells (INS1E cell line and rat islets) and enteroendocrine L cells (56, 57). Nevertheless, the mechanisms of sustained alterations in AKT and ERK phosphorylation were not fully understood. In our *in vitro* model, AKT and ERK phosphorylation was suppressed at all time points during 30-min acute insulin stimulation, suggesting that the insulin resistance we observed was impaired responsiveness - consistent with signaling deficiencies at both the receptor level and in post-receptor components (58). Indeed, multiple components of insulin signaling were reduced at the transcriptional level revealed by our transcriptomics analysis. Our observations also verified the distinct responses to hyperinsulinemia on the bifurcate insulin signaling pathways. Chronic 200 nM insulin treatment preferentially increased basal AKT phosphorylation - as a sign of sustained activation - but did not increase the basal ERK phosphorylation, possibly due to desensitization, as reported in neurons (59). Diet- and hyperinsulinemia-induced insulin resistance is generally considered to be related specifically to AKT phosphorylation. Chemical inhibition of the AKT pathway using the non-selective PI3K inhibitor LY-294002, but not ERK pathway inhibition, has been reported to protect insulin resistance both *in vitro* and *in vivo* (59, 60). In addition to the downregulation of INSR protein abundance, we found a subtle increase in INSR internalization, and future studies will be required to determine the importance and mechanisms of this phenomenon. Further work is also required to understand the interplay between INSR expression and both major branches of downstream signaling.

A major observation of our work is that *Insr* mRNA was directly reduced by hyperinsulinemia in cultured cells, consistent with reports from other cell culture systems (46, 47), and also is consistently negatively correlated with fasting insulin in both mouse and human skeletal muscle *in vivo*. Indeed, T2D patients were found to have lower *Insr* mRNA expression in skeletal muscle biopsies (61). Notably, hyperglycemia can increase *Insr* expression in lymphocyte and cancer cell lines (62, 63), while high glucose inhibits β-cell *Insr* expression through autocrine insulin action and INSR-FOXO1 signaling (62, 63). Interestingly, glucose only induces insulin resistance in the presence of insulin in cultured hepatocytes, adipocytes and skeletal muscle (64-66). Therefore, reduced *Insr* expression by hyperinsulinemia may be a key, independent factor of INSR downregulation and insulin resistance.

Intermittent fasting, time-restricted feeding, caloric restriction, and carbohydrate restriction positively modify risk factors in diabetes, including reducing hyperinsulinemia, increasing insulin sensitivity, improving β-cell responsiveness, and lowering the levels of circulating glucose (67-69). Several human trials suggest that fasting regimes can be more effective for reducing insulin and increasing insulin sensitivity than they are for reducing glucose (70, 71). By mimicking the low-insulin state, the serum starvation phase of our studies revealed some possible molecular mechanisms of the beneficial effects of fasting on muscle cells, including the restoration of protein phosphorylation in insulin signaling pathways and partial recovery of *Insr* transcription, INSR protein and overall transcriptomic changes. These data hint that some deleterious effects of hyperinsulinemia are reversible but may require a long enough period of reduced insulin exposure to ‘reset’.

Pathway analysis of the transcriptomics correctly revealed broad effects of hyperinsulinemia and serum starvation on insulin signaling and FOXO signaling pathways and highlighted potential upstream transcription factors. This further supports utility of robust RNA pathway analysis alone (Stokes et al 2020) to correctly identify key protein regulators in muscle tissue (72). Besides FOXO1, other transcription factors such as SP1, HMGA1, C/EBPβ and NUCKS have been reported to regulate *Insr* expression (73-75). We identified at least one novel transcriptional repressor of the *Insr* gene, SIN3A, which was upregulated by hyperinsulinemia. SIN3A interacts with histone deacetylases, typically HDAC1/2, to inhibit transcription and interacts with other transcription factors that were identified in our informatics analyses (76). For example, SIN3A and MYC inhibit each other and form a negative feedback loop (77). MAX dimerizes with either MYC or MXI1 (MAD family protein) in a competing manner to activate or repress target genes (78), and MXI1 recruits SIN3A for gene inhibition (78). Interestingly, the knockdown level we achieved for MYC, MAX, MXI1 did not have significant effects on the transcription of *Insr* in muscle. One possibility is that the roles of these transcription factors on *Insr* are indirect and rely on the action of SIN3A, while another possibility is that SIN3A acts through alternative pathways. A recent study identified SIN3A as a FOXO1 corepressor of the glucokinase gene in the liver (79) and this may represent a possible sight of interaction in our system.

Despite its inherent reductionism, our *in vitro* model identified plausible molecular features underpinning the descriptive relationship between hyperinsulinemia in the development of insulin resistance and T2D. The trancriptomic responses in our muscle cell model were reflective of the correlation analysis of human skeletal muscle, indicating that it represents an informative resource to interrogate hyperinsulinemia-induced insulin resistance. We demonstrated that *in vitro* hyperinsulinemia and serum ‘fasting’ have profound effects on AKT and ERK signaling, INSR abundance and localization, and transcriptional activities. Future additional characterization of the effect of hyperinsulinemism on INSR trafficking, degradation, and detailed post-receptor alterations on protein level will provide a greater understanding of the role of hyperinsulinemia in promoting obesity (9) and diabetes.

## Acknowledgments

We acknowledge Dr. Brain Rodrigues for providing the C2C12 cells and Dr. Stephane Flibotte in LSI Bioinformatics Core for advice. We thank our colleagues Xiaoke Hu and Leanne Beet for performing the insulin radioimmunoassays. We thank those who provided helpful comments on previous drafts of this manuscript posted on *Biorxiv*.

## Supplementary figure legends

**Figure S1. Insulin dose- and time-depend acute signaling in the hyperinsulinemia-induced insulin resistance model**. Myotubes cultured in control (0 nM insulin) or hyperinsulinemic (200 nM insulin) medium were stimulated with acute 0.2, 2 or 20 nM insulin for 1, 5, 10, 15 or 30 min after serum starvation. **(A)** phospho-AKT (T308, S473), **(B)** phospho-ERK1/2, and **(C)** INSR phosphorylation were measured. (n=4; ^#^ effect of hyperinsulinemia, ^&^ effect of acute insulin, × interaction between two factors, Mixed Effect Model.)

**Figure S2. RNA-seq analysis of hyperinsulinemia and serum starvation highlighting glucose metabolism and FOXO signaling pathways. (A)** Top 50 most significantly altered genes with lowest p value. **(B)** Selected KEGG pathways enriched from genes downregulated by hyperinsulinemia before

starvation (BS, 0 vs 200 nM insulin). **(C)** Upregulated genes (BS, 200 vs 0 nM) enriched under Reactome pathway “Glucose metabolism”. **(D)** Heatmap showing relative expression levels of genes under “Glucose metabolism” and “Glucolysis” Reactome pathways. Unmarked genes were differentially expressed by hyperinsulinemia only before starvation (BS, 200 vs 0 nM) but not after starvation. Genes marked by * were differentially expressed both before and after starvation (BS & AS, 200 vs 0 nM). Genes marked by # were only altered by starvation (200 nM, AS vs BS) but not by hyperinsulinemia. **(E)** The log2 fold change of the common of DE genes before (BS, 200 vs 0 nM) and after starvation (AS, 200 vs 0 nM).

**Figure S3. Mouse and human skeletal muscle data sets. (A)** Common DE genes in mouse IRMOE model and our hyperinsulinemia model (BS, 200 vs 0 nM). **(B)** Common DE genes in mouse hyperinsulinemic clamp study and our hyperinsulinemia model (BS, 200 vs 0 nM). **(C)** Fasting insulin levels of subjects in the 3 human data sets.

**Figure S4. Endocytosis-related differentially expressed genes and surface biotinylation assay. (A)** Differentially expressed genes related to endocytosis pathways. **(B)** Scheme of surface biotinylation assay to measure surface or internalized INSR. Surface or internalized INSR in protein lysates were detected by western blots.

**Table S1**. Differentially expressed genes by *in vitro* hyperinsulinemia and starvation

**Table S2**. Reactome pathway enrichment from differentially expressed genes

**Table S3**. KEGG pathway enrichment from differentially expressed genes

**Table S4**. Comparison of differentially expressed genes in our *in vitro* model, IRMOE and clamp studies

**Table S5**. Comparison of insulin-correlated genes in SMP and FUSION human skeletal muscle and differentially expressed genes in our *in vitro* model

## Notes

### Competing Interest Statement

The authors have declared no competing interest.

### Summary of Updates

This version of the manuscript includes RNA sequencing results and analysis of human transcriptomes.

## References

1. Defronzo, R. A. (2009) Banting Lecture. From the triumvirate to the ominous octet: a new paradigm for the treatment of type 2 diabetes mellitus. Diabetes 58, 773–795

2. Prentki, M., and Nolan, C. J. (2006) Islet beta cell failure in type 2 diabetes. J Clin Invest 116, 1802–1812

3. Corkey, B. E. (2012) Banting lecture 2011: hyperinsulinemia: cause or consequence? Diabetes 61, 4–13

4. Shanik, M. H., Xu, Y., Skrha, J., Dankner, R., Zick, Y., and Roth, J. (2008) Insulin resistance and hyperinsulinemia: is hyperinsulinemia the cart or the horse? Diabetes Care 31 Suppl 2, S262–268

5. Page, M. M., and Johnson, J. D. (2018) Mild Suppression of Hyperinsulinemia to Treat Obesity and Insulin Resistance. Trends Endocrinol Metab 29, 389–399

6. Le Stunff, C., and Bougneres, P. (1994) Early changes in postprandial insulin secretion, not in insulin sensitivity, characterize juvenile obesity. Diabetes 43, 696–702

7. Spadaro, L., Alagona, C., Palermo, F., Piro, S., Calanna, S., Parrinello, G., Purrello, F., and Rabuazzo, A. M. (2011) Early phase insulin secretion is increased in subjects with normal fasting glucose and metabolic syndrome: a premature feature of beta-cell dysfunction. Nutr Metab Cardiovasc Dis 21, 206–212

8. Trico, D., Natali, A., Arslanian, S., Mari, A., and Ferrannini, E. (2018) Identification, pathophysiology, and clinical implications of primary insulin hypersecretion in nondiabetic adults and adolescents. JCI Insight 3

9. Wiebe, N., Ye, F., Crumley, E. T., Bello, A., Stenvinkel, P., and Tonelli, M. (2021) Temporal Associations Among Body Mass Index, Fasting Insulin, and Systemic Inflammation: A Systematic Review and Meta-analysis. JAMA Netw Open 4, e211263

10. Dankner, R., Chetrit, A., Shanik, M. H., Raz, I., and Roth, J. (2012) Basal state hyperinsulinemia in healthy normoglycemic adults heralds dysglycemia after more than two decades of follow up. Diabetes Metab Res Rev 28, 618–624

11. Zimmet, P. Z., Collins, V. R., Dowse, G. K., and Knight, L. T. (1992) Hyperinsulinaemia in youth is a predictor of type 2 (non-insulin-dependent) diabetes mellitus. Diabetologia 35, 534–541

12. Templeman, N. M., Flibotte, S., Chik, J. H. L., Sinha, S., Lim, G. E., Foster, L. J., Nislow, C., and Johnson, J. D. (2017) Reduced Circulating Insulin Enhances Insulin Sensitivity in Old Mice and Extends Lifespan. Cell Rep 20, 451–463

13. Mehran, A. E., Templeman, N. M., Brigidi, G. S., Lim, G. E., Chu, K. Y., Hu, X., Botezelli, J. D., Asadi, A., Hoffman, B. G., Kieffer, T. J., Bamji, S. X., Clee, S. M., and Johnson, J. D. (2012) Hyperinsulinemia drives diet-induced obesity independently of brain insulin production. Cell Metab 16, 723–737

14. Page, M. M., Skovso, S., Cen, H., Chiu, A. P., Dionne, D. A., Hutchinson, D. F., Lim, G. E., Szabat, M., Flibotte, S., Sinha, S., Nislow, C., Rodrigues, B., and Johnson, J. D. (2018) Reducing insulin via conditional partial gene ablation in adults reverses diet-induced weight gain. FASEB J 32, 1196–1206

15. Hamza, S. M., Sung, M. M., Gao, F., Soltys, C. M., Smith, N. P., MacDonald, P. E., Light, P. E., and Dyck, J. R. (2017) Chronic insulin infusion induces reversible glucose intolerance in lean rats yet ameliorates glucose intolerance in obese rats. Biochim Biophys Acta Gen Subj 1861, 313–322

16. Yang, X., Mei, S., Gu, H., Guo, H., Zha, L., Cai, J., Li, X., Liu, Z., and Cao, W. (2014) Exposure to excess insulin (glargine) induces type 2 diabetes mellitus in mice fed on a chow diet. J Endocrinol 221, 469–480

17. Marangou, A. G., Weber, K. M., Boston, R. C., Aitken, P. M., Heggie, J. C., Kirsner, R. L., Best, J. D., and Alford, F. P. (1986) Metabolic consequences of prolonged hyperinsulinemia in humans. Evidence for induction of insulin insensitivity. Diabetes 35, 1383–1389

18. Del Prato, S., Leonetti, F., Simonson, D. C., Sheehan, P., Matsuda, M., and DeFronzo, R. A. (1994) Effect of sustained physiologic hyperinsulinaemia and hyperglycaemia on insulin secretion and insulin sensitivity in man. Diabetologia 37, 1025–1035

19. Gregory, J. M., Smith, T. J., Slaughter, J. C., Mason, H. R., Hughey, C. C., Smith, M. S., Kandasamy, B., Greeley, S. A. W., Philipson, L. H., Naylor, R. N., Letourneau, L. R., Abumrad, N. N., Cherrington, A. D., and Moore, D. J. (2019) Iatrogenic Hyperinsulinemia, Not Hyperglycemia, Drives Insulin Resistance in Type 1 Diabetes as Revealed by Comparison to GCK-MODY (MODY2). Diabetes

20. Rome, S., Clément, K., Rabasa-Lhoret, R., Loizon, E., Poitou, C., Barsh, G. S., Riou, J. P., Laville, M., and Vidal, H. (2003) Microarray profiling of human skeletal muscle reveals that insulin regulates approximately 800 genes during a hyperinsulinemic clamp. J Biol Chem 278, 18063–18068

21. O’Brien, R. M., and Granner, D. K. (1996) Regulation of gene expression by insulin. Physiol Rev 76, 1109–1161

22. Batista, T. M., Garcia-Martin, R., Cai, W., Konishi, M., O’Neill, B. T., Sakaguchi, M., Kim, J. H., Jung, D. Y., Kim, J. K., and Kahn, C. R. (2019) Multi-dimensional Transcriptional Remodeling by Physiological Insulin In Vivo. Cell Rep 26, 3429-3443.e3423

23. Di Camillo, B., Irving, B. A., Schimke, J., Sanavia, T., Toffolo, G., Cobelli, C., and Nair, K. S. (2012) Function-based discovery of significant transcriptional temporal patterns in insulin stimulated muscle cells. PLoS One 7, e32391

24. Tseng, L. T., Lin, C. L., Tzen, K. Y., Chang, S. C., and Chang, M. F. (2013) LMBD1 protein serves as a specific adaptor for insulin receptor internalization. J Biol Chem 288, 32424–32432

25. Rowzee, A. M., Ludwig, D. L., and Wood, T. L. (2009) Insulin-like growth factor type 1 receptor and insulin receptor isoform expression and signaling in mammary epithelial cells. Endocrinology 150, 3611–3619

26. Harith, H. H., Di Bartolo, B. A., Cartland, S. P., Genner, S., and Kavurma, M. M. (2016) Insulin promotes vascular smooth muscle cell proliferation and apoptosis via differential regulation of tumor necrosis factor-related apoptosis-inducing ligand. J Diabetes 8, 568–578

27. Zhang, J., Tang, H., Zhang, Y., Deng, R., Shao, L., Liu, Y., Li, F., Wang, X., and Zhou, L. (2014) Identification of suitable reference genes for quantitative RT-PCR during 3T3-L1 adipocyte differentiation. Int J Mol Med 33, 1209–1218

28. Dobin, A., Davis, C. A., Schlesinger, F., Drenkow, J., Zaleski, C., Jha, S., Batut, P., Chaisson, M., and Gingeras, T. R. (2013) STAR: ultrafast universal RNA-seq aligner. Bioinformatics 29, 15–21

29. Love, M. I., Huber, W., and Anders, S. (2014) Moderated estimation of fold change and dispersion for RNA-seq data with DESeq2. Genome Biol 15, 550

30. Wang, G., Yu, Y., Cai, W., Batista, T. M., Suk, S., Noh, H. L., Hirshman, M., Nigro, P., Li, M. E., Softic, S., Goodyear, L., Kim, J. K., and Kahn, C. R. (2020) Muscle-Specific Insulin Receptor Overexpression Protects Mice From Diet-Induced Glucose Intolerance but Leads to Postreceptor Insulin Resistance. Diabetes 69, 2294–2309

31. Yu, G., Wang, L. G., Han, Y., and He, Q. Y. (2012) clusterProfiler: an R package for comparing biological themes among gene clusters. OMICS 16, 284–287

32. Yu, G., and He, Q. Y. (2016) ReactomePA: an R/Bioconductor package for reactome pathway analysis and visualization. Mol Biosyst 12, 477–479

33. Zhou, G., Soufan, O., Ewald, J., Hancock, R. E. W., Basu, N., and Xia, J. (2019) NetworkAnalyst 3.0: a visual analytics platform for comprehensive gene expression profiling and meta-analysis. Nucleic Acids Res 47, W234–W241

34. Timmons, J. A., Atherton, P. J., Larsson, O., Sood, S., Blokhin, I. O., Brogan, R. J., Volmar, C. H., Josse, A. R., Slentz, C., Wahlestedt, C., Phillips, S. M., Phillips, B. E., Gallagher, I. J., and Kraus, W. E. (2018) A coding and non-coding transcriptomic perspective on the genomics of human metabolic disease. Nucleic Acids Res 46, 7772–7792

35. Welsh, E. A., Eschrich, S. A., Berglund, A. E., and Fenstermacher, D. A. (2013) Iterative rank-order normalization of gene expression microarray data. BMC Bioinformatics 14, 153

36. Møller, A. B., Kampmann, U., Hedegaard, J., Thorsen, K., Nordentoft, I., Vendelbo, M. H., Møller, N., and Jessen, N. (2017) Altered gene expression and repressed markers of autophagy in skeletal muscle of insulin resistant patients with type 2 diabetes. Sci Rep 7, 43775

37. Kampmann, U., Christensen, B., Nielsen, T. S., Pedersen, S. B., Ørskov, L., Lund, S., Møller, N., and Jessen, N. (2011) GLUT4 and UBC9 protein expression is reduced in muscle from type 2 diabetic patients with severe insulin resistance. PLoS One 6, e27854

38. Harrison, X. A., Donaldson, L., Correa-Cano, M. E., Evans, J., Fisher, D. N., Goodwin, C. E. D., Robinson, B. S., Hodgson, D. J., and Inger, R. (2018) A brief introduction to mixed effects modelling and multi-model inference in ecology. PeerJ 6, e4794

39. Bates, D., Machler, M., Bolker, B. M., and Walker, S. C. (2015) Fitting Linear Mixed-Effects Models Using lme4. J Stat Softw 67, 1–48

40. Melmed, S., Polonsky, K. S., Larsen, P. R., and Kronenberg, H. M. (2017) Williams Textbook of Endocrinology, Elsevier Saunders

41. McAuley, K. A., Williams, S. M., Mann, J. I., Walker, R. J., Lewis-Barned, N. J., Temple, L. A., and Duncan, A. W. (2001) Diagnosing insulin resistance in the general population. Diabetes Care 24, 460–464

42. Boucher, J., Kleinridders, A., and Kahn, C. R. (2014) Insulin receptor signaling in normal and insulin-resistant states. Cold Spring Harb Perspect Biol 6

43. Fröjdö, S., Vidal, H., and Pirola, L. (2009) Alterations of insulin signaling in type 2 diabetes: a review of the current evidence from humans. Biochim Biophys Acta 1792, 83–92

44. Morikawa, Y., Ueyama, E., and Senba, E. (2004) Fasting-induced activation of mitogen-activated protein kinases (ERK/p38) in the mouse hypothalamus. J Neuroendocrinol 16, 105–112

45. Taylor, D. L., Jackson, A. U., Narisu, N., Hemani, G., Erdos, M. R., Chines, P. S., Swift, A., Idol, J., Didion, J. P., Welch, R. P., Kinnunen, L., Saramies, J., Lakka, T. A., Laakso, M., Tuomilehto, J., Parker, S. C. J., Koistinen, H. A., Davey Smith, G., Boehnke, M., Scott, L. J., Birney, E., and Collins, F. S. (2019) Integrative analysis of gene expression, DNA methylation, physiological traits, and genetic variation in human skeletal muscle. Proc Natl Acad Sci U S A 116, 10883–10888

46. Okabayashi, Y., Maddux, B. A., McDonald, A. R., Logsdon, C. D., Williams, J. A., and Goldfine, I. D. (1989) Mechanisms of insulin-induced insulin-receptor downregulation. Decrease of receptor biosynthesis and mRNA levels. Diabetes 38, 182–187

47. Zhang, Z., Li, X., Liu, G., Gao, L., Guo, C., Kong, T., Wang, H., Gao, R., Wang, Z., and Zhu, X. (2011) High insulin concentrations repress insulin receptor gene expression in calf hepatocytes cultured in vitro. Cell Physiol Biochem 27, 637–640

48. Sbraccia, P., D’Adamo, M., Leonetti, F., Caiola, S., Iozzo, P., Giaccari, A., Buongiorno, A., and Tamburrano, G. (1996) Chronic primary hyperinsulinaemia is associated with altered insulin receptor mRNA splicing in muscle of patients with insulinoma. Diabetologia 39, 220–225

49. Orengo, D. J., Aguadé, M., and Juan, E. (2017) Characterization of dFOXO binding sites upstream of the Insulin Receptor P2 promoter across the Drosophila phylogeny. Plos One 12, e0188357

50. Puig, O., and Tjian, R. (2005) Transcriptional feedback control of insulin receptor by dFOXO/FOXO1. Genes Dev 19, 2435–2446

51. Ni, Y. G., Wang, N., Cao, D. J., Sachan, N., Morris, D. J., Gerard, R. D., Kuro-O, M., Rothermel, B. A., and Hill, J. A. (2007) FoxO transcription factors activate Akt and attenuate insulin signaling in heart by inhibiting protein phosphatases. Proc Natl Acad Sci U S A 104, 20517–20522

52. Nakae, J., Kitamura, T., Ogawa, W., Kasuga, M., and Accili, D. (2001) Insulin regulation of gene expression through the forkhead transcription factor Foxo1 (Fkhr) requires kinases distinct from Akt. Biochemistry 40, 11768–11776

53. Jochen, A. L., Berhanu, P., and Olefsky, J. M. (1986) Insulin Internalization and Degradation in Adipocytes from Normal and Type-Ii Diabetic Subjects. J Clin Endocr Metab 62, 268–274

54. Trischitta, V., Gullo, D., Squatrito, S., Pezzino, V., Goldfine, I. D., and Vigneri, R. (1986) Insulin Internalization into Monocytes Is Decreased in Patients with Type-Ii Diabetes-Mellitus. J Clin Endocr Metab 62, 522–528

55. El Akoum, S., Lamontagne, V., Cloutier, I., and Tanguay, J. F. (2011) Nature of fatty acids in high fat diets differentially delineates obesity-linked metabolic syndrome components in male and female C57BL/6J mice. Diabetol Metab Syndr 3

56. Rachdaoui, N., Polo-Parada, L., and Ismail-Beigi, F. (2019) Prolonged Exposure to Insulin Inactivates Akt and Erk1/2 and Increases Pancreatic Islet and INS1E beta-Cell Apoptosis. J Endocr Soc 3, 69–90

57. Lim, G. E., Huang, G. J., Flora, N., LeRoith, D., Rhodes, C. J., and Brubaker, P. L. (2009) Insulin regulates glucagon-like peptide-1 secretion from the enteroendocrine L cell. Endocrinology 150, 580–591

58. Kahn, C. R. (1978) Insulin resistance, insulin insensitivity, and insulin unresponsiveness: a necessary distinction. Metabolism 27, 1893–1902

59. Kim, B., McLean, L. L., Philip, S. S., and Feldman, E. L. (2011) Hyperinsulinemia induces insulin resistance in dorsal root ganglion neurons. Endocrinology 152, 3638–3647

60. Liu, H. Y., Hong, T., Wen, G. B., Han, J., Zuo, D., Liu, Z., and Cao, W. (2009) Increased basal level of Akt-dependent insulin signaling may be responsible for the development of insulin resistance. Am J Physiol Endocrinol Metab 297, E898–906

61. Palsgaard, J., Brøns, C., Friedrichsen, M., Dominguez, H., Jensen, M., Storgaard, H., Spohr, C., Torp-Pedersen, C., Borup, R., De Meyts, P., and Vaag, A. (2009) Gene expression in skeletal muscle biopsies from people with type 2 diabetes and relatives: differential regulation of insulin signaling pathways. Plos One 4, e6575

62. Martinez, S. C., Cras-Méneur, C., Bernal-Mizrachi, E., and Permutt, M. A. (2006) Glucose regulates Foxo1 through insulin receptor signaling in the pancreatic islet beta-cell. Diabetes 55, 1581–1591

63. Briata, P., Briata, L., and Gherzi, R. (1990) Glucose starvation and glycosylation inhibitors reduce insulin receptor gene expression: characterization and potential mechanism in human cells. Biochem Biophys Res Commun 169, 397–405

64. Liu, H. Y., Cao, S. Y., Hong, T., Han, J., Liu, Z., and Cao, W. (2009) Insulin is a stronger inducer of insulin resistance than hyperglycemia in mice with type 1 diabetes mellitus (T1DM). J Biol Chem 284, 27090–27100

65. Garvey, W. T., Olefsky, J. M., and Marshall, S. (1985) Insulin receptor down-regulation is linked to an insulin-induced postreceptor defect in the glucose transport system in rat adipocytes. J Clin Invest 76, 22–30

66. Ciaraldi, T. P., Abrams, L., Nikoulina, S., Mudaliar, S., and Henry, R. R. (1995) Glucose transport in cultured human skeletal muscle cells. Regulation by insulin and glucose in nondiabetic and non-insulin-dependent diabetes mellitus subjects. J Clin Invest 96, 2820–2827

67. Mattson, M. P., Longo, V. D., and Harvie, M. (2017) Impact of intermittent fasting on health and disease processes. Ageing Res Rev 39, 46–58

68. Patterson, R. E., and Sears, D. D. (2017) Metabolic Effects of Intermittent Fasting. Annu Rev Nutr 37, 371–393

69. Nuttall, F. Q., Almokayyad, R. M., and Gannon, M. C. (2015) Comparison of a carbohydrate-free diet vs. fasting on plasma glucose, insulin and glucagon in type 2 diabetes. Metabolism 64, 253–262

70. Sutton, E. F., Beyl, R., Early, K. S., Cefalu, W. T., Ravussin, E., and Peterson, C. M. (2018) Early Time-Restricted Feeding Improves Insulin Sensitivity, Blood Pressure, and Oxidative Stress Even without Weight Loss in Men with Prediabetes. Cell Metab 27, 1212–1221 e1213

71. Williams, K. V., Mullen, M. L., Kelley, D. E., and Wing, R. R. (1998) The effect of short periods of caloric restriction on weight loss and glycemic control in type 2 diabetes. Diabetes Care 21, 2–8

72. Stokes, T., Timmons, J. A., Crossland, H., Tripp, T. R., Murphy, K., McGlory, C., Mitchell, C. J., Oikawa, S. Y., Morton, R. W., Phillips, B. E., Baker, S. K., Atherton, P. J., Wahlestedt, C., and Phillips, S. M. (2020) Molecular Transducers of Human Skeletal Muscle Remodeling under Different Loading States. Cell Rep 32, 107980

73. Foti, D., Iuliano, R., Chiefari, E., and Brunetti, A. (2003) A nucleoprotein complex containing Sp1, C/EBP beta, and HMGI-Y controls human insulin receptor gene transcription. Mol Cell Biol 23, 2720–2732

74. Foti, D., Chiefari, E., Fedele, M., Iuliano, R., Brunetti, L., Paonessa, F., Manfioletti, G., Barbetti, F., Brunetti, A., Croce, C. M., and Fusco, A. (2005) Lack of the architectural factor HMGA1 causes insulin resistance and diabetes in humans and mice. Nat Med 11, 765–773

75. Qiu, B., Shi, X., Wong, E. T., Lim, J., Bezzi, M., Low, D., Zhou, Q., Akıncılar, S. C., Lakshmanan, M., Swa, H. L., Tham, J. M., Gunaratne, J., Cheng, K. K., Hong, W., Lam, K. S., Ikawa, M., Guccione, E., Xu, A., Han, W., and Tergaonkar, V. (2014) NUCKS is a positive transcriptional regulator of insulin signaling. Cell Rep 7, 1876–1886

76. Kadamb, R., Mittal, S., Bansal, N., Batra, H., and Saluja, D. (2013) Sin3: insight into its transcription regulatory functions. Eur J Cell Biol 92, 237–246

77. Nascimento, E. M., Cox, C. L., MacArthur, S., Hussain, S., Trotter, M., Blanco, S., Suraj, M., Nichols, J., Kübler, B., Benitah, S. A., Hendrich, B., Odom, D. T., and Frye, M. (2011) The opposing transcriptional functions of Sin3a and c-Myc are required to maintain tissue homeostasis. Nat Cell Biol 13, 1395–1405

78. Zervos, A. S., Gyuris, J., and Brent, R. (1993) Mxi1, a protein that specifically interacts with Max to bind Myc-Max recognition sites. Cell 72, 223–232

79. Langlet, F., Haeusler, R. A., Lindén, D., Ericson, E., Norris, T., Johansson, A., Cook, J. R., Aizawa, K., Wang, L., Buettner, C., and Accili, D. (2017) Selective Inhibition of FOXO1 Activator/Repressor Balance Modulates Hepatic Glucose Handling. Cell 171, 824-835.e818

