## Supplemental Figures for "Transcriptomic analysis of human and mouse muscle during hyperinsulinemia demonstrates insulin receptor downregulation as a mechanism for insulin resistance"

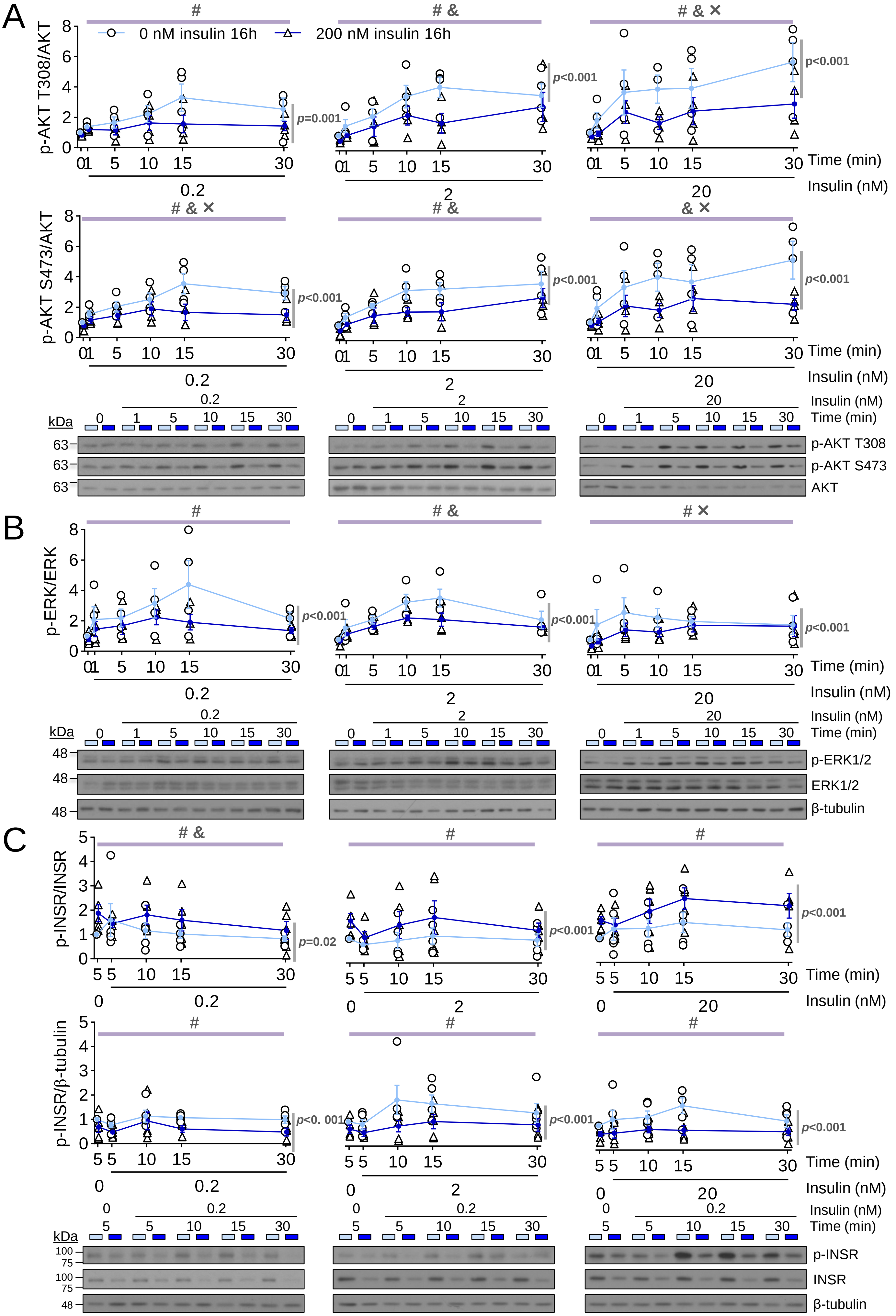


**Figure S1. Insulin dose- and time-depend acute signaling in the hyperinsulinemia-induced insulin resistance model.**

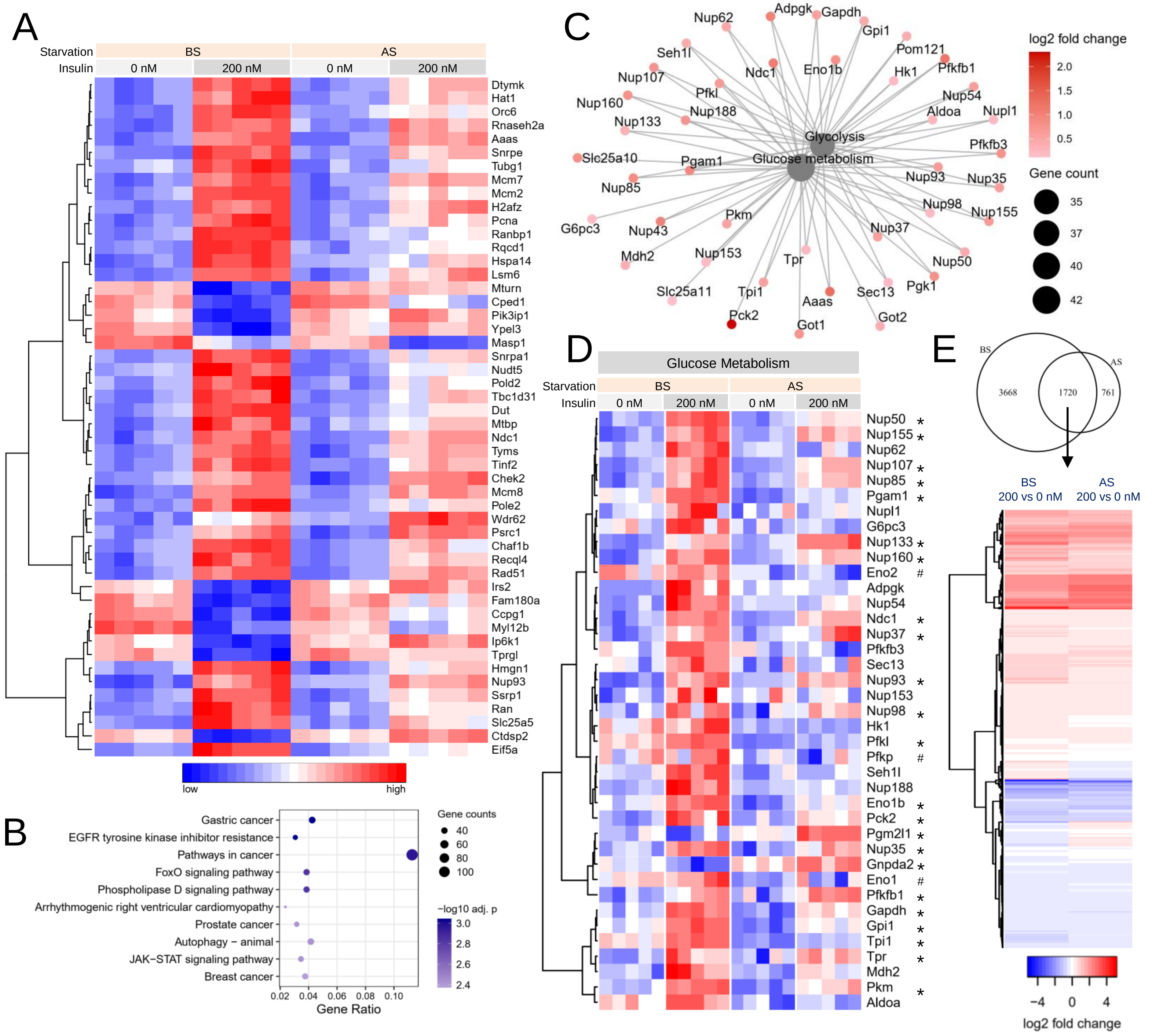


**Figure S2. RNA-seq analysis of hyperinsulinemia and serum starvation highlighting glucose metabolism and FOXO signaling pathways.**

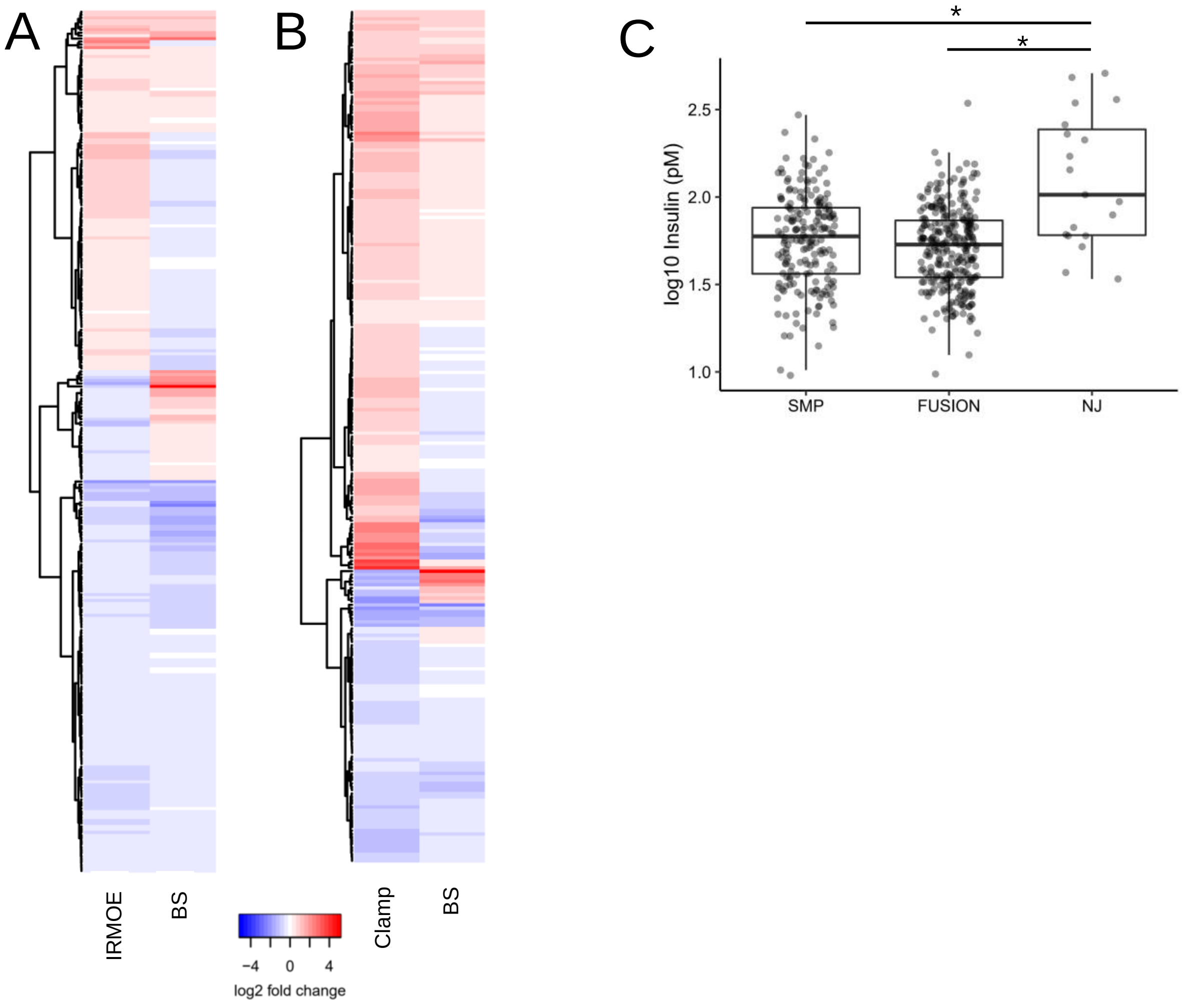


**Figure S3. Mouse and human skeletal muscle data sets.**

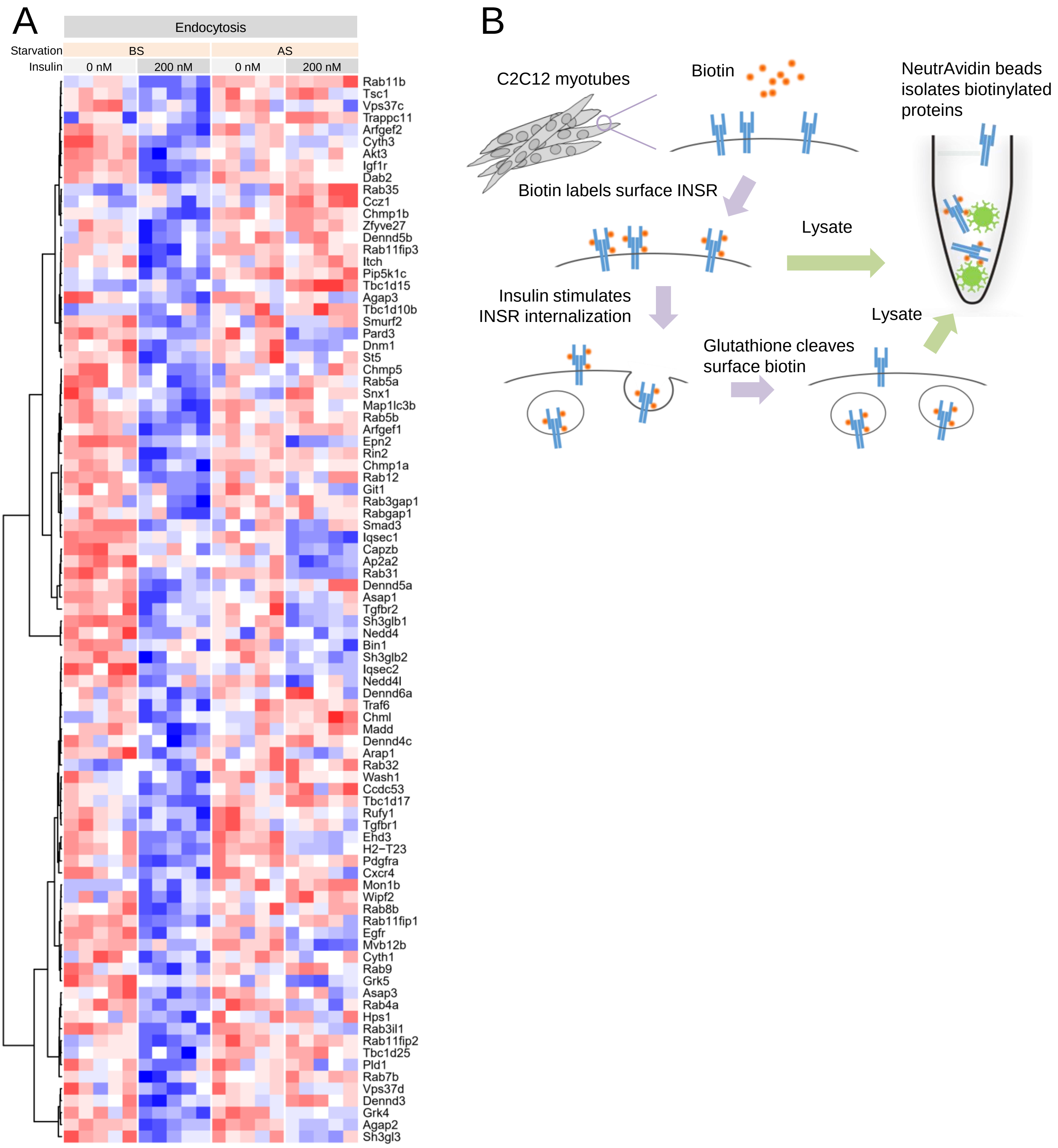


**Figure S4. Endocytosis related differentially expressed genes and surface biotinylation assay.**
